# Microbial exoenzymes catalyzed the transition to an oxygenated Earth

**DOI:** 10.1101/2025.09.11.675588

**Authors:** Alia Sanger, Andrew D. Steen, Joanne S. Boden, Stella Cellier-Goetghebeur, Maria Bayder, Elliott P. Mueller, Galen P. Halverson, Nicolas B. Cowan, Rika E. Anderson, Joelle N. Pelletier, Eva E. Stüeken, Mojtaba Fakhraee, Kurt O. Konhauser, Nagissa Mahmoudi

## Abstract

Microbial exoenzymes—extracellular enzymes secreted to degrade complex organic polymers— are essential for recycling carbon and nutrients, thus sustaining primary productivity in today’s oceans^1^. Yet, their evolutionary history and role in shaping Earth’s early biosphere remain entirely unexplored. Here, we trace the origins of microbial exoenzymes and reveal their previously unrecognized role in driving planetary oxygenation. Our results show that exoenzymes are more common in microorganisms utilizing high-energy metabolisms, likely reflecting the energetic costs of enzyme biosynthesis and secretion. They are especially advantageous in environments rich in particulate organic matter (POM). A refined carbon cycle model indicates that early Archean oceans offered few such habitats, as low productivity and intense UV radiation rapidly photodegraded POM. However, with a Paleoproterozoic rise of atmospheric oxygen^2^, increased oxidative weathering boosted marine primary productivity and POM accumulation^3^, creating conditions favoring exoenzyme evolution. Molecular clock analyses further indicate that alkaline phosphatase, a key phosphorus-releasing exoenzyme, had likely emerged with the permanent rise of oxygen, enabling more efficient phosphorus recycling. We propose that exoenzymes initiated a positive feedback loop: by accelerating nutrient regeneration, they fueled cyanobacterial productivity and oxygen release, which in turn favored greater exoenzyme capacity, reinforcing long-term oxygenation of the planet.

## Main

Microorganisms are central to the cycling of carbon, nitrogen, and phosphorus on Earth through their roles in both primary production of biomass and the remineralization of organic matter in the oceans and sediments. While low-molecular-weight organic matter, typically < 1000 Da (i.e., dissolved organic matter, DOM) can be directly assimilated by cells, a substantial fraction of organic matter exists as structurally complex, high-molecular weight polymers, known as particulate organic matter (POM) that are too large to be taken up by cells. The degradation of this recalcitrant material is primarily mediated by heterotrophic microorganisms that secrete exoenzymes to hydrolyze POM into low-molecular-weight DOM that can be transported across cell membranes^4^. By enabling access to otherwise inaccessible organic polymers, exoenzymes regulate the efficiency of microbial recycling and exert a key influence on the fate of bioessential elements in the marine environment^1^.

Exoenzymes represent a critical but underexplored driver of Earth’s geochemical and biological evolution^5^. The metabolic ability to degrade POM was essential for regenerating and redistributing critical nutrients and metals in early oceans^6,7^. In addition, the efficiency of heterotrophic recycling directly influences the burial rates of POM in marine sediments, a key sink for reduced carbon. Both processes —nutrient regeneration^8,9^ and POM burial^10^ —have been recognized as fundamental controls on planetary oxygenation. The permanent accumulation of oxygen in Earth’s atmosphere, termed the Great Oxidation Event (GOE), occurred broadly between ca. 2.5 and 2.2 billion years ago^2^, irreversibly altering Earth’s surface environments and biosphere. Despite extension studies of exoenzymes in modern oceans^11,12^, their evolutionary history and potential contribution to Earth’s oxygenation remain unresolved.

In this work, we investigate the evolutionary trajectory of exoenzymes and their role in POM cycling and nutrient dynamics through Earth’s history. First, we quantify the genomic distribution of exoenzymes across microbial metabolisms to assess how changing environmental redox conditions shaped their prevalence in ancient oceans. Second, we analyze the sequence diversity of predicted exoenzymes to evaluate whether homologous enzymes are shared across metabolic groups, implying common function and ancestry. Third, to explore the ecological context that favoured exoenzyme evolution, we apply a biogeochemical model of the marine carbon cycle, focusing on how ocean chemistry and primary productivity influenced POM-associated habitats. Finally, we integrate gene tree–species tree reconciliations with molecular clock analyses to constrain the timing of key exoenzyme innovations, specifically those involved in the hydrolysis of marine organic phosphorus compounds.

### Exoenzymes are concentrated in microbial metabolisms with higher energetic potentials

Heterotrophic microorganisms oxidize organic matter by energetically coupling its degradation to the reduction of terminal electron acceptors, such as oxygen, nitrate, ferric iron, and sulfate. However, it remains unclear how the genes responsible for encoding secreted exoenzymes— critical for degrading POM—are conserved across metabolic groups. Resolving this question is critical for understanding the evolutionary trajectory of exoenzymes and their role in shaping carbon and nutrient cycling in early marine environments. To address this, we quantified the distribution of exoenzymes in extant microbial genomes representing distinct metabolic groups. Here, metabolic groups are defined as microorganisms that share a common strategy for obtaining energy and nutrients, such as reliance on specific electron acceptors or light, regardless of their taxonomic classification. We compiled a dataset of 3,076 high-quality bacterial and archaeal genomes representing both cultured and uncultured microorganisms, using key marker genes as a proxy for metabolic groups (**Table S1, S2**). We screened these genomes for 26 known classes of exoenzymes targeting proteins and carbohydrates (**Table S3**), which together comprise the bulk of bioavailable marine organic matter. Hydrolytic enzymes may be localized in the cytoplasm, periplasm or associated with the cell wall, or may be secreted freely into the extracellular medium. They can also be involved in essential intracellular functions such as degrading misfolded or damaged proteins (e.g. peptidases). While hydrolytic enzymes themselves are ubiquitous, the export of hydrolytic enzymes to the external environment (i.e. exoenzymes) represents a distinct foraging strategy for acquiring carbon and nutrients from extracellular POM. We identified putative exoenzymes based on multiple criteria, including the presence of signal peptide sequences and distinctive signatures in secondary structure and amino acid composition (see Supplemental Materials). This approach allowed us to quantify the fraction of genomes within each metabolic group that likely possess the capacity to produce exoenzymes.

Our analysis reveals that microbial exoenzymes are disproportionately concentrated within a small subset of metabolisms. Strikingly, 93% of all putative exoenzymes are encoded by genomes associated with just four metabolic groups: dissimilatory ferric iron reduction (31%), aerobic heterotrophy (23%), dissimilatory nitrate reduction (22%), and fermentation (17%) (**Fig. 1**). By contrast, genomes associated with dissimilatory sulfate reduction and methanogenesis encoded few, if any, detectable exoenzymes. While the role of fermenters in the initial degradation of POM in anoxic environments is well established, our results extend the genetic capacity for exoenzyme production to other anaerobic metabolic groups, namely ferric iron and nitrate reducers. This finding is consistent with observations from modern sediments, where degradation of POM is enhanced under nitrate-rich^13^ or ferric iron-rich conditions^14^. Conversely, the paucity of exoenzyme genes in sulfate-reducing and methanogenic microorganisms supports radiotracer experiments in Baltic Sea sediments, which show steady POM degradation rates across sulfate- and methane-rich zones, indicating that neither of these pathways controls the initial, rate-limiting hydrolytic step^15^. Collectively, our findings identify the key microbial players involved in POM cycling across redox gradients and establish a mechanistic framework for interpreting organic matter turnover in both modern and ancient environments.

**Figure 1.**
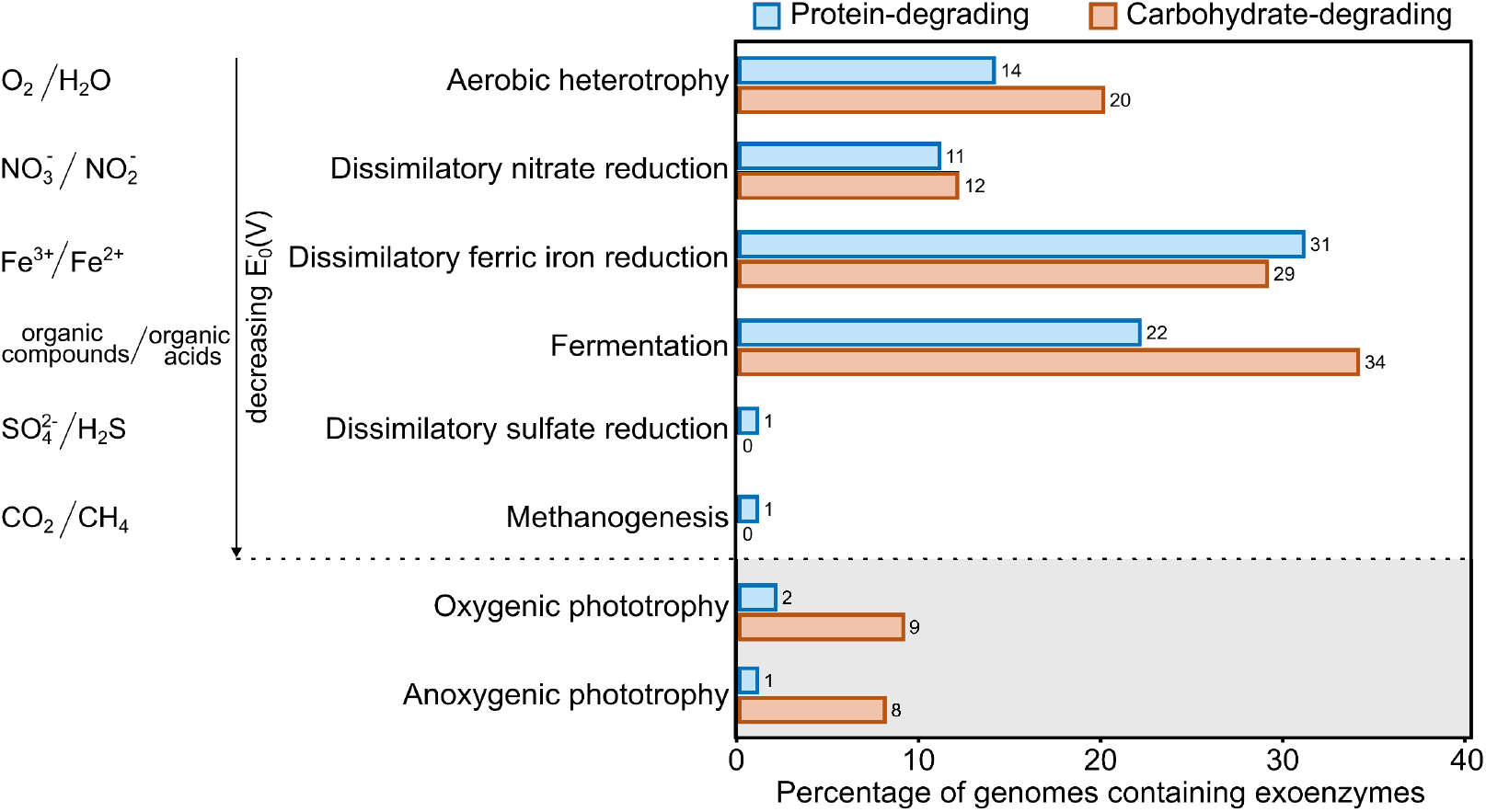
Exoenzymes are concentrated in metabolic groups with higher energetic potential. The percentage of genomes that contain at least one predicted protein-degrading (blue) or carbohydrate-degrading (orange) exoenzyme across major metabolic groups. Heterotrophic metabolisms are arranged by the standard reduction potentials (E_0_′) in volts (V) of their respective redox reactions, reflecting the energy yield available from coupling the oxidation of organic compounds to the reduction of a terminal electron acceptor. A larger E_0_′ difference between donor and acceptor corresponds to a greater release of free energy available to the cell. Electron acceptors with the highest oxidizing potential, and thus greatest energy yield, appear at the top of the redox hierarchy. The relative E_0_′ are based on standard conditions (25 °C, pH 7, 1 atm). Broad phototrophic groups are shown separately, as they do not rely solely on thermodynamic differences between redox pairs; instead, they use light energy to drive electron excitation and transfer.

The uneven distribution of exoenzymes across metabolic groups likely reflects the fundamental constraints imposed by cellular energetics and the high costs of enzyme production. The synthesis and secretion of exoenzymes requires considerable investments of carbon, nitrogen, and energy^16^. Moreover, since microorganisms lack mechanisms to re-import proteins once secreted, exoenzymes are effectively lost to the environment^17^, placing strong selective pressure on cells to tightly regulate their production^18^. Evolution appears to have favoured strategies that mitigate these costs: extracellular proteins often contain disproportionately few energetically costly amino acids^19^, and aquatic exopeptidases preferentially liberate energetically valuable amino acids^20^. These patterns point to a broader evolutionary logic, whereby exoenzyme deployment is fine-tuned to the metabolic capabilities of the host and the energetic returns associated with substrate degradation. In this framework, microbial lineages appear to optimize their hydrolytic investment in response to both substrate availability and the thermodynamic efficiency of their energy metabolism.

Here we find that exoenzyme production is largely restricted to metabolisms that generate higher free energy and ATP molecules through respiration using energetically favorable terminal electron acceptors (**Fig. 1**). A notable exception is fermentation, which operates independently of exogenous electron acceptors and instead generates energy via substrate-level phosphorylation^21^. Although fermentation yields less free energy and ATP per reaction, it is characterized by lower minimum catabolic power requirements and higher anabolic biomass yields per mole of substrate compared to other anaerobic pathways^22,23^. This enhanced metabolic efficiency may allow fermenters to allocate a larger portion of their energy budget toward exoenzyme production, conferring this hydrolytic capability to an otherwise energetically “weak” metabolism. Together, these findings support the idea that exoenzyme production is governed by an energetic threshold, whereby only those metabolisms capable of meeting the bioenergetic costs can effectively access POM via extracellular hydrolysis.

To further explore hydrolytic capabilities across metabolic groups, we analyzed 1,307 predicted exoenzyme sequences using sequence similarity networks (SSNs) (**Fig. 2**). These SSNs revealed that sequences cluster primarily by enzyme functional class, forming distinct groups. In contrast, there is little evidence of clustering by metabolic groups because closely related exoenzymes are distributed across phylogenetically and metabolically diverse microorganisms. This pattern suggests that microorganisms with different core metabolisms can have similar hydrolytic capabilities. It further supports the notion that some microorganisms function as “generalist” exoenzyme producers, capable of secreting a broad repertoire of exoenzymes^24^. Previous work examining the genetic potential of three specific exoenzymes showed their presence across all major prokaryotic phyla, although this capacity was often restricted to smaller, closely related clades or ecotypes^12^. This uneven distribution of exoenzyme potential among heterotrophic microorganisms likely reflects an evolutionary response to ecological pressures, such as substrate or habitat availability, with natural selection favouring this trait in specific clades that benefit from extracellular hydrolysis.

**Figure 2.**
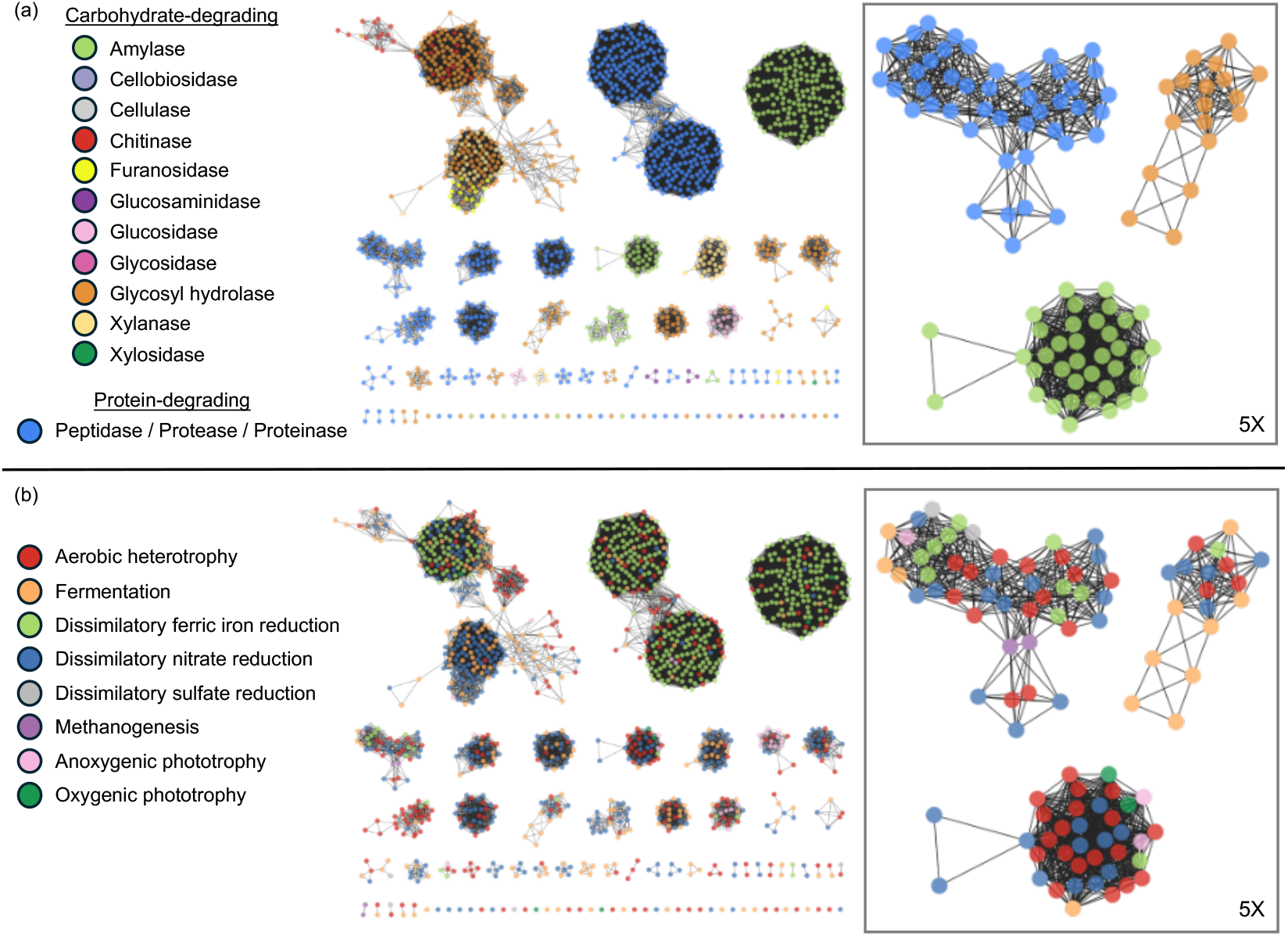
Diverse metabolisms share functionally and genetically similar exoenzymes. Sequence similarity network of predicted exoenzymes associated with (A) enzyme type based on sequence annotation; and (B) metabolic guild. The 1307 exoenzyme protein sequences are depicted by individual nodes (colored circles). Edges (grey lines) between the nodes indicate a pairwise BLAST e-value of 1×10^-30^. Each exoenzyme sequence is represented by a node (colored circle); 1,267 nodes are grouped into 48 clusters (e-value of 1×10^-30^) based on high sequence similarity, though 40 sequences were dissimilar from all other sequences and are thus not connected to any other nodes. Featured clusters (right panels) depict how diverse metabolic guilds share functionally and genetically similar exoenzymes.

### Ecological selection for exoenzymes in particle-rich oceans

In the modern ocean, exoenzymes facilitate the stepwise breakdown of POM into high- and then low-molecular-weight DOM, ultimately enabling assimilation into microbial biomass. Planktonic microorganisms drift freely in the ocean and consume DOM, while others grow by attaching to and degrading POM. Within organic aggregates, both exoenzymes and their hydrolysis products are more efficiently retained in the surrounding matrix^17,25^. This retention makes exoenzymes particularly advantageous in diffusion-limited settings, such as marine particles, where they provide benefits for particle-attached microorganisms. Consequently, the availability of POM likely acted as an important ecological and evolutionary driver, expanding the range of habitats in which exoenzyme production conferred a selective advantage.

To investigate how particulate habitat availability may have changed through Earth’s history, we developed a biogeochemical model that simulates the major sources and sinks of dissolved organic carbon (DOC) and infers implications for particulate organic carbon (POC) in the ocean. DOC and POC are reliable indicators for tracking large-scale dynamics of DOM and POM since they comprise the largest fractions of these reservoirs and are quantified using standardized methods across global datasets^26^. Our approach builds on a previously established framework that reconstructs the evolution of marine carbon reservoirs over geological timescales^27^, with refined parameterization to account for the effects of high ultraviolet (UV) radiation fluxes that reached Earth’s surface prior to the formation of a protective ozone layer^28,29^. Under these high-UV conditions, enhanced photochemical reactions would have accelerated the degradation of POM into DOM, thereby reducing the abundance of particle-associated habitats, such as water column particles or bottom sediment, to heterotrophic microorganisms^30–32^. A full description of the model, including parameterization of early Earth UV fluxes and photochemical processes, is provided in the Supplementary Materials.

Our model implies that long-term, secular changes in ocean chemistry reshaped the ecological pressures influencing exoenzyme production over geological time (**Fig. 3, top panel**). Simulated DOC concentrations remain relatively stable, reflecting a persistent reservoir of labile carbon in the water column, consistent with empirical records^33^. By contrast, primary productivity—and with it, POC availability—likely increased markedly through time, though not explicitly simulated, consistent with independent estimates of rising productivity^34^. In the Archean ocean, primary productivity may have been just 0.1%-1% of modern levels, limiting the formation of particle-rich habitats. In parallel, high UV fluxes would have accelerated the photochemical conversion of POM into DOM, further reducing the prevalence of habitats that favor exoenzyme production. Moreover, the relative abundance of DOM per microbial cell would have been much greater than in modern oceans, implying that early heterotrophs inhabited a DOM-rich ecosystem with weak selective pressures for exoenzyme production. This chemical and ecological landscape would have supported a small, low-energy biosphere dominated by free-living heterotrophs with limited access to POM.

**Figure 3.**
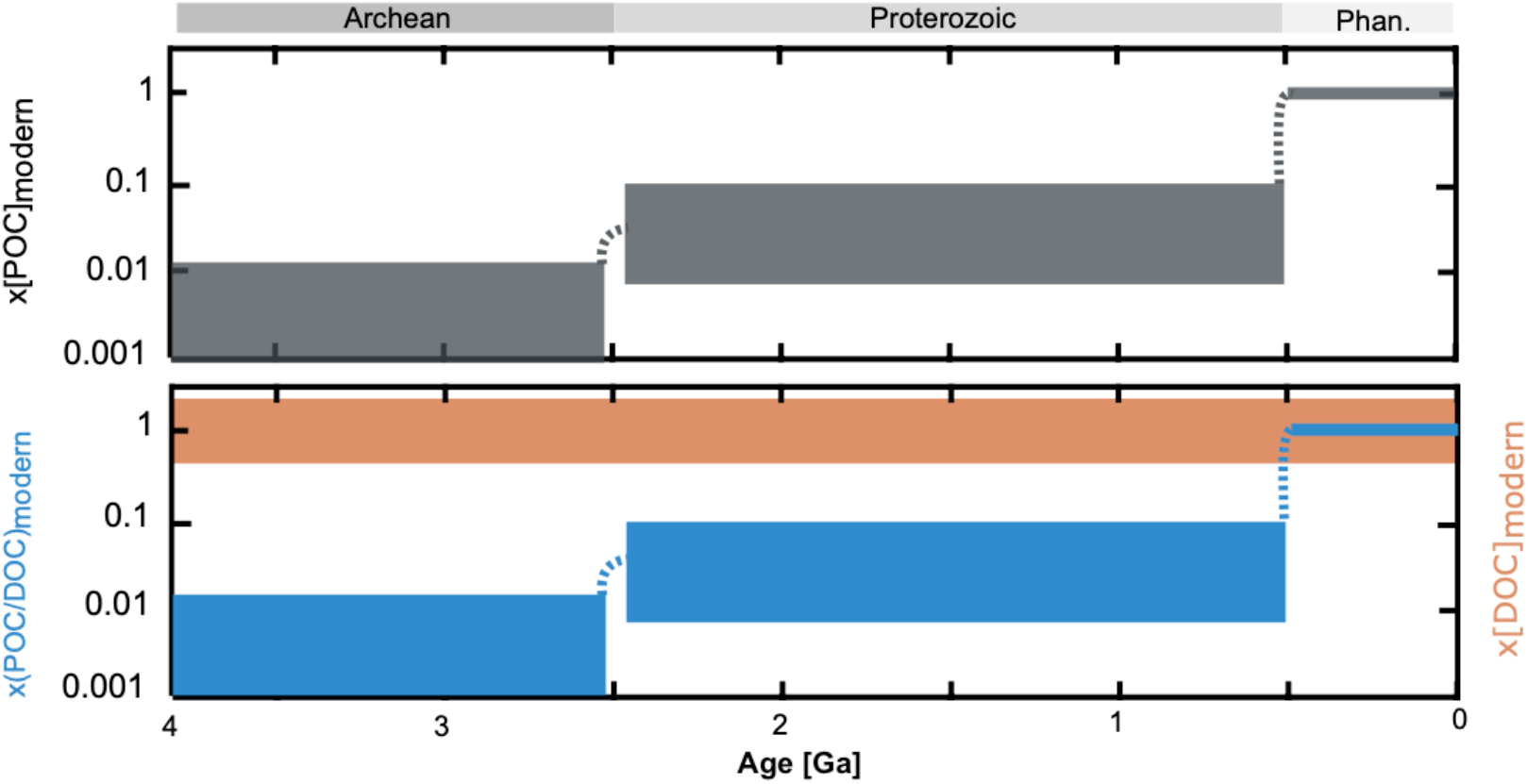
Relative reservoir sizes of particulate organic carbon (POC) and dissolved organic carbon (DOC) over Earth history, normalized to modern values. The top panel shows the evolution of the POC reservoir size relative to the modern, while the bottom panel shows the evolution of the POC/DOC ratio (blue, left axis) and the DOC reservoir size relative to modern (orange, right axis). Geological eras (Archean, Proterozoic, Phanerozoic) are indicated at the top. Note the logarithmic scale on the y-axes.

The GOE marked a pivotal shift in the balance between dissolved and particulate pools of organic matter in the oceans (**Fig. 3, bottom panel**). The rise of an ozone layer in the early Paleoproterozoic, perhaps as early as ∼2.4 billion years ago^35^, though possibly moderated by iodine emissions^36^, reduced UV-driven photodegradation of POM. Coupled with increasing primary productivity and microbial cell densities^34^, this atmospheric shift likely expanded the marine POM reservoir and created more particle-rich habitats where exoenzyme production presented a selective advantage. In this ecological context, higher POC-to-DOC ratios and greater cell densities would have intensified selective pressures to degrade more complex particulate substrates. At the same time, rising cell densities would have favored cooperative behaviors such as POM degradation, which becomes more efficient under crowded conditions^37^. Experimental and modeling studies support this interpretation, showing that efficient POM turnover by particle-attached microbes only occurs once a critical cell density is reached^38^. Together, these ecological and chemical transformations following the GOE promoted the proliferation of exoenzymes, facilitating broader microbial access to POM, and shaping overall organic matter cycling throughout the Proterozoic.

To examine when specific exoenzymes likely emerged and spread on the early Earth, we conducted gene tree-species tree reconciliations in conjunction with molecular clock analyses to estimate the timing of the rise and spread of extracellular alkaline phosphatases across the tree of life. These enzymes catalyze the hydrolysis of organic phosphorus compounds, allowing microorganisms to more efficiently scavenge phosphorus^39–41^. We focused on extracellular alkaline phosphatases for two key reasons. First, they are substantially more conserved in both gene sequence and substrate specificity than other exoenzymes, such as extracellular peptidases, making them more suitable for molecular clock estimates. Second, phosphorus has long been considered a limiting nutrient for primary productivity throughout Earth’s history^42^, making alkaline phosphatases important mediators of phosphorus recycling in ancient oceans.

We identified putative alkaline phosphatase sequences from a curated tree of life comprising 865 genomes^43^ (see Supplemental Materials). Gene sequences were aligned to construct gene trees, which were then reconciled with the corresponding species tree to infer the timing of alkaline phosphatase speciation, duplication, or transfer events. Because alkaline phosphatases can function both within and outside of the cell, we focused our analysis on monophyletic gene clades for which we were confident that the ancestral enzymes were extracellular (**Fig. S1**). Our analyses show that extracellular alkaline phosphatases within these clades likely emerged sometime between the Neoarchean to Paleoproterozoic (**Tables S6-S9**). Although molecular clock estimates carry inherent uncertainties, this timing is broadly consistent with robust signals of oxygen appearing and rising on Earth, supporting the hypothesis that the evolutionary expansion of exoenzymes—and with them, enhanced microbial phosphorus regeneration—was a key feature of the early Proterozoic oceans.

### Co-evolution of exoenzymes and Earth’s surface redox state

Our findings reveal a dynamic interplay between microbial metabolism and ocean redox conditions, suggesting that redox state has not only shaped the distribution of microbial metabolisms but also exerted a fundamental influence on carbon burial efficiency and nutrient regeneration over geological time. The rate of carbon burial depends on both the net primary productivity (NPP) and the efficiency with which that POM is remineralized. We demonstrate that the genetic capacity to produce exoenzymes is disproportionately concentrated amongst aerobic heterotrophs, dissimilatory ferric iron and nitrate reducers, and fermenters. This distribution implies that the evolution and ecological prominence of these metabolisms directly impacted organic matter degradation and nutrient recycling, and thereby Earth’s long-term carbon cycle (**Fig. 4**). As such, microbial control over biogeochemical fluxes has been intimately coupled to the planet’s evolving redox landscape.

**Figure 4.**
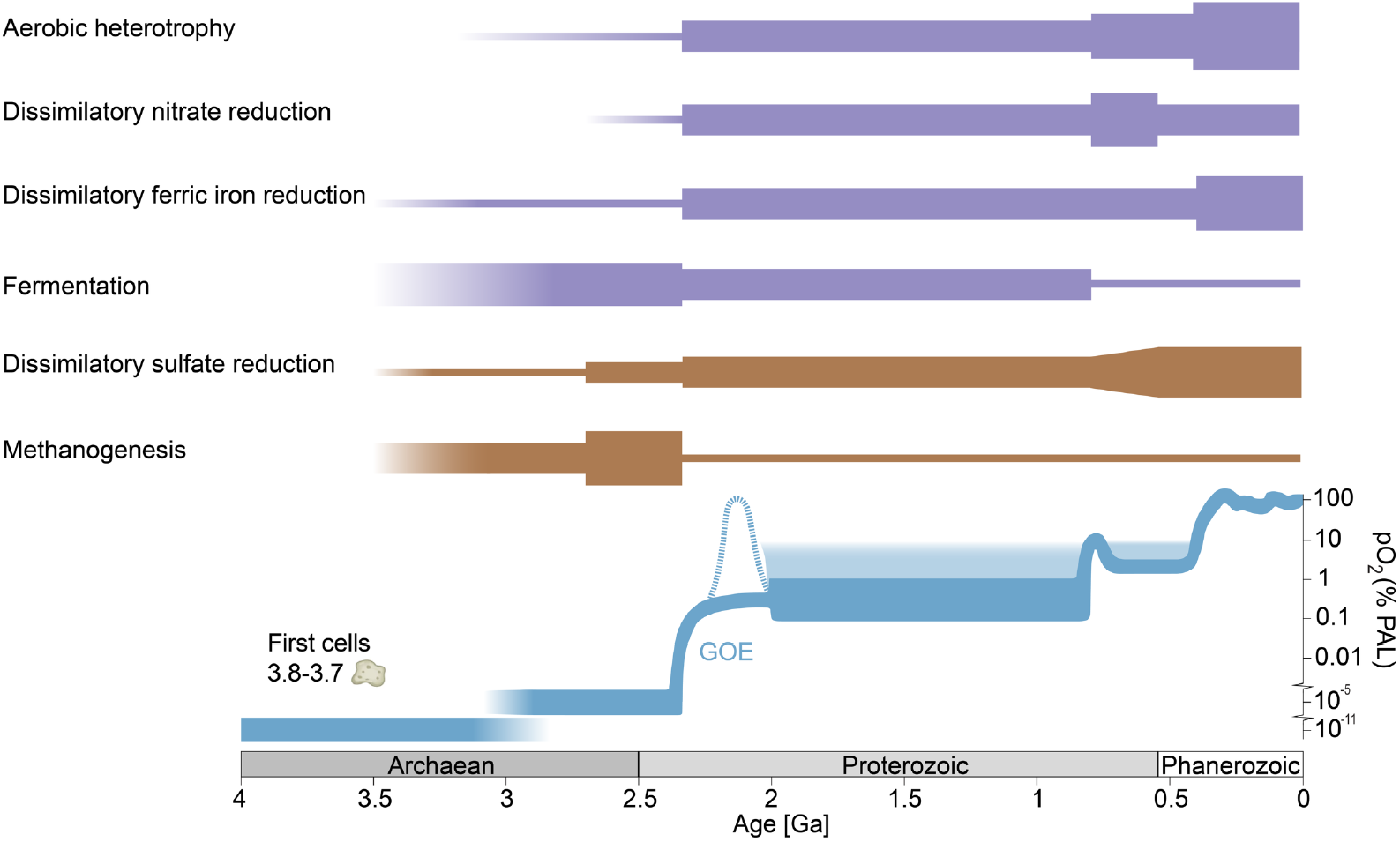
Relative emergence and ecological impact of heterotrophic metabolisms through time. The emergence of each metabolism is inferred from a combination of geochemical, isotopic and phylogenetic evidence summarized in Lyons et al. (2024)^51^. Line thickness represents the relative ecological impact of each metabolic group, ranging from a baseline (thinnest) to peak influence (thickest). Shifts in ecological impact, whether expansion or contraction, are broadly associated with changes in the oxidation state of the biosphere. Metabolic groups with abundant exoenzymes are shaded purple; those with few exoenzymes are shaded brown. The blue curve (bottom, adapted from Lyons et al. 2021)^59^ depicts the evolution of atmospheric oxygen throughout Earth history relative to the present atmospheric level (PAL). The solid blue curve denotes states with considerable geochemical evidence whereas the dashed and graded sections of the blue curve indicate less certain states and transitions.

Both phylogenetic analyses^44,45^ and geochemical evidence^46,47^ suggest that aerobic heterotrophs and dissimilatory nitrate reducers emerged just prior to the GOE, in the later Neoarchean, thus constraining the ecological expansion of these groups until after the GOE. Dissimilatory ferric iron reduction also emerged prior to the GOE^48^ and may have been widespread on the outer continental shelf, as indicated by the prevalence of reduced diagenetic minerals in banded iron formations^49^. However, this metabolism was probably restricted to upwelling zones where ferric iron was available^50^. Dissimilatory ferric iron reduction is thought to have greatly expanded in the early Paleoproterozoic, as rising oxygen levels oxidized Fe^2+^ to Fe^3+^, increasing the supply of ferric iron for microbial reduction^51^. Taken together, these patterns suggest that prior to the GOE, the recycling of POM in the Archean oceans could have hinged on the metabolic activity of fermenters (**Fig. 5**). This reliance on a low-energy metabolism would have resulted in lower organic matter remineralization rates, thereby limiting the release of nutrients back into the water column and increasing the proportion of biomass from primary productivity that escaped degradation^52^.

**Figure 5.**
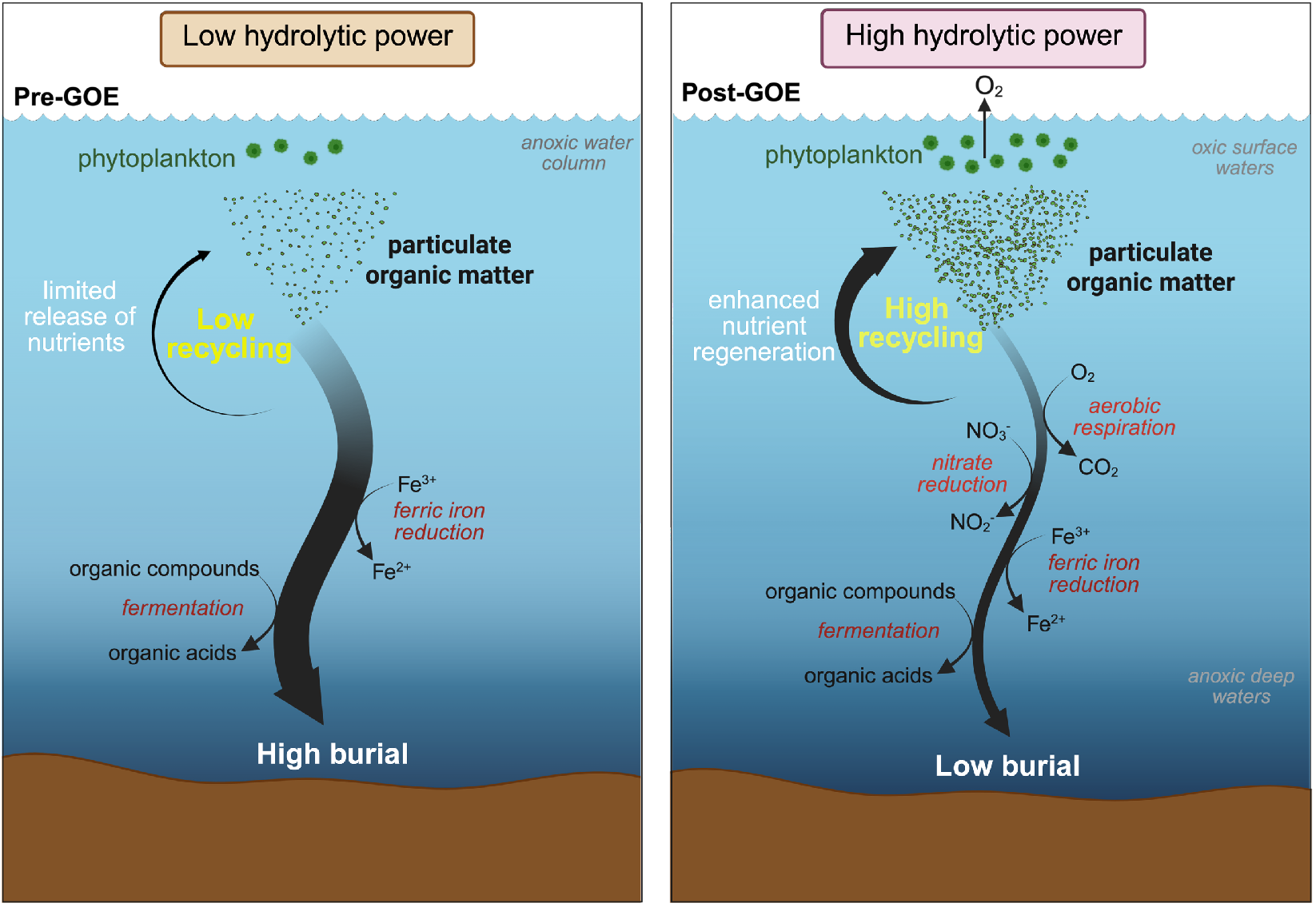
Conceptual model for the evolution of exoenzymes and the recycling of organic matter across the Great Oxidation Event (GOE). (A) In the Archean Ocean, prior to the GOE, the recycling of organic matter depended entirely on the activity of fermenters. This resulted in low recycling efficiency, elevated carbon burial, and limited regeneration of essential nutrients back into seawater. This scarcity of particulate organic matter (POM) also reduced the availability of niches for microbial colonization, thus limiting the selective pressure for the evolution of exoenzyme systems. (b) Following the GOE, rising oxygen levels increased the availability of high-energy electron acceptors (e.g. O_2_, nitrate and ferric iron), supporting the expansion of metabolic groups with high exoenzyme capacity, such as aerobic heterotrophs, dissimilatory nitrate reducers and dissimilatory ferric iron reducers. Concurrently, increases in POM abundance and microbial cell densities would have amplified the selective pressure to produce exoenzymes. Collectively, these changes in the Paleoproterozoic promoted more efficient recycling of organic matter and regeneration of nutrients back into the water column, ultimately fueling greater primary productivity and oxygen release. This feedback between exoenzyme evolution, nutrient cycling, and oxygen production would have amplified Earth’s progressive oxygenation.

This interpretation is consistent with both the sedimentary rock record and geochemical estimates of primary productivity during the Archean^34^. Although primary productivity was substantially lower than today, total organic carbon (TOC) concentrations in Archean sedimentary rocks are comparable to those in younger Proterozoic and Phanerozoic strata (**Fig. S2**). We propose that this apparent contradiction can be reconciled by the inefficient microbial recycling of POM due to limited exoenzyme production, which allowed a greater fraction of POM to escape degradation and become buried in the sediment pile. This would have enhanced carbon burial and increased the proportion of primary producer biomass preserved in sediments. Indeed, previous estimates suggest Archean burial efficiencies ranged from a few percent up to tens of percent, far exceeding the modern ocean average of 0.2%^53,54^. Moreover, recent findings have independently proposed that suppressed organic matter remineralization is required to account for high organic matter content in some Neoarchean shales^55^. While suppressed remineralization of organic matter in Archean oceans has traditionally been attributed to the lack of terminal electron acceptors for heterotrophic respiration such as oxygen and sulfate^52^, our results reveal that a more fundamental constraint lay in the scarcity of efficient exoenzymatic degradation pathways (**Fig. 5**). Inefficient microbial recycling of organic matter would not only have enhanced carbon burial but also restricted the release of bioessential nutrients (e.g. phosphorus and trace metals) back into the water column, thereby constraining primary productivity and possibly delaying the ecological expansion of oxygenic photoautotrophs^8,9,42,56–58^.

Our results identify exoenzymes as a pivotal innovation that drove marine organic matter turnover and nutrient cycling, emerging in concert with energetic and ecological transformations in the wake of the GOE. During the Archean, low primary productivity, intense UV-driven photodegradation, and a paucity of particle-associated habitats suppressed selective pressure for exoenzyme production. Early heterotrophic communities lived in a DOM-rich, low-energy biosphere with limited POM, while inefficient recycling of key nutrients constrained the growth of primary producers. The global rise of oxygen during the Paleoproterozoic marked a turning point: greater primary productivity, reduced UV fluxes, and increasing microbial cell densities shifted seawater toward higher POM-to-DOM ratios, expanding the ecological niches where exoenzyme production conferred a competitive advantage. Rising cell densities also fostered cooperative behaviors such as extracellular hydrolysis that enhanced POM degradation, while the emergence of more energetically favorable terminal electron acceptors enabled the evolution of metabolic strategies capable of meeting the substantial energetic costs of exoenzyme production. Collectively, this led to more efficient degradation of POM. Our molecular clock analysis suggests that extracellular alkaline phosphatase, a key enzyme for phosphorus recycling, arose during this transition, reinforcing the coupling between POM degradation, nutrient recycling, and rising productivity in Paleoproterozoic oceans.

In conclusion, these intertwined changes established a positive feedback loop between primary producers and heterotrophic consumers. The rise of cyanobacteria increased oxygen levels and POM availability through increased microbial biomass, promoting the emergence of exoenzyme-producing heterotrophs. In turn, enhanced nutrient recycling sustained higher primary productivity and further oxygen release. This self-reinforcing feedback reshaped global carbon and nutrient cycles, driving the progressive oxygenation of Earth’s surface environments. Recognizing this coupling between microbial innovation and ecological interactions offers a new perspective on the co-evolution of life and planetary habitability.

## ACKNOWLEDGEMENTS

This work was supported by the Heising-Simons Foundation (grant no. 2023-4653 to NM and NBC) and New Frontiers in Research Fund Exploration grant (grant no. NFRFE-2019-00794 to NM and JP). EES and JSB acknowledge funding from a NERC Frontiers grant (NE/V010824/1 awarded to EES). ADS acknowledges funding from NSF OCE-2145434. REA was supported as part of the Virtual Planetary Laboratory Team, which is a member of the NASA Nexus for Exoplanet System Science, and funded via the NASA Astrobiology Program ICAR Grant 80NSSC231398.

## AUTHOR CONTRIBUTIONS

AS: study design, data acquisition and analysis, writing-editing; ADS: conceptualization, study design, data analysis, writing-editing; JB: data acquisition and analysis, writing-editing; RA: data analysis, writing-editing; SCG: data analysis; MB: data analysis; EM: writing-editing; GPH: writing-editing; NBC: data acquisition and analysis; JNP: data analysis, writing-editing; EES: data acquisition and analysis, writing-editing; MF: conceptualization, data acquisition and analysis, writing-editing; KOK: conceptualization, writing-editing; NM: conceptualization, study design, data analysis, writing-editing, funding, supervision.

## DATA AND CODE AVAILABILITY

The data used for all analyses presented in this article along with supporting code for data analysis can be found at https://figshare.com/s/5304713d4e372cdfd593.

## REFERENCES

1. Arnosti, C. Microbial Extracellular Enzymes and the Marine Carbon Cycle. Annu. Rev. Mar. Sci. 3, 401–425 (2011).

2. Poulton, S. W. et al. A 200-million-year delay in permanent atmospheric oxygenation. Nature 592, 232–236 (2021).

3. Konhauser, K. O. et al. Aerobic bacterial pyrite oxidation and acid rock drainage during the Great Oxidation Event. Nature 478, 369–373 (2011).

4. Benz, R. & Bauer, K. Permeation of hydrophilic molecules through the outer membrane of gram‐negative bacteria: Review of bacterial porins. Eur. J. Biochem. 176, 1–19 (1988).

5. Mahmoudi, N., Steen, A. D., Halverson, G. P. & Konhauser, K. O. Biogeochemistry of Earth before exoenzymes. Nat. Geosci. 16, 845–850 (2023).

6. Anbar, A. D. Elements and Evolution. Science 322, 1481–1483 (2008).

7. Robbins, L. J. et al. Trace elements at the intersection of marine biological and geochemical evolution. Earth-Sci. Rev. 163, 323–348 (2016).

8. Alcott, L. J., Mills, B. J. W. & Poulton, S. W. Stepwise Earth oxygenation is an inherent property of global biogeochemical cycling. Science 366, 1333–1337 (2019).

9. Alcott, L. J., Mills, B. J. W., Bekker, A. & Poulton, S. W. Earth’s Great Oxidation Event facilitated by the rise of sedimentary phosphorus recycling. Nat. Geosci. 15, 210–215 (2022).

10. Holland, H. D. The Chemistry of the Atmosphere and Oceans. (Wiley, New York, 1978).

11. Zhao, Z., Baltar, F. & Herndl, G. J. Linking extracellular enzymes to phylogeny indicates a predominantly particle-associated lifestyle of deep-sea prokaryotes. Sci. Adv. 6, eaaz4354 (2020).

12. Zimmerman, A. E., Martiny, A. C. & Allison, S. D. Microdiversity of extracellular enzyme genes among sequenced prokaryotic genomes. ISME J. 7, 1187–1199 (2013).

13. Bulseco, A. N. et al. Nitrate addition stimulates microbial decomposition of organic matter in salt marsh sediments. Glob. Change Biol. 25, 3224–3241 (2019).

14. Aller, R. C., Mackin, J. E. & Cox, R. T. Diagenesis of Fe and S in Amazon inner shelf muds: apparent dominance of Fe reduction and implications for the genesis of ironstones. Cont. Shelf Res. 6, 263–289 (1986).

15. Beulig, F., Røy, H., Glombitza, C. & Jørgensen, B. B. Control on rate and pathway of anaerobic organic carbon degradation in the seabed. Proc. Natl. Acad. Sci. 115, 367–372 (2018).

16. Allison, S. D., Weintraub, M. N., Gartner, T. B. & Waldrop, M. P. Evolutionary-Economic Principles as Regulators of Soil Enzyme Production and Ecosystem Function. in Soil Enzymology (eds. Shukla, G. & Varma, A.) vol. 22 229–243 (Springer Berlin Heidelberg, Berlin, Heidelberg, 2010).

17. Vetter, Y. A., Deming, J. W., Jumars, P. A. & Krieger-Brockett, B. B. A Predictive Model of Bacterial Foraging by Means of Freely Released Extracellular Enzymes. Microb. Ecol. 36, 75–92 (1998).

18. Allison, S. D. Cheaters, diffusion and nutrients constrain decomposition by microbial enzymes in spatially structured environments. Ecol. Lett. 8, 626–635 (2005).

19. Smith, D. R. & Chapman, M. R. Economical Evolution: Microbes Reduce the Synthetic Cost of Extracellular Proteins. mBio 1, e00131–10 (2010).

20. Steen, A. D., Vazin, J. P., Hagen, S. M., Mulligan, K. H. & Wilhelm, S. W. Substrate specificity of aquatic extracellular peptidases assessed by competitive inhibition assays using synthetic substrates. Aquat. Microb. Ecol. 75, 271–281 (2015).

21. White, D. The Physiology and Biochemistry of Prokaryotes. (Oxford University Press, New York, 1995).

22. Jaussi, M. et al. Cell-specific rates of sulfate reduction and fermentation in the sub-seafloor biosphere. Front. Microbiol. 14, 1198664 (2023).

23. LaRowe, D. E. & Amend, J. P. Power limits for microbial life. Front. Microbiol. 6, (2015).

24. Newton, R. J. et al. Genome characteristics of a generalist marine bacterial lineage. ISME J. 4, 784–798 (2010).

25. Smith, D. C., Simon, M., Alldredge, A. L. & Azam, F. Intense hydrolytic enzyme activity on marine aggregates and implications for rapid particle dissolution. Nature 359, 139–142 (1992).

26. Liang, Z., Letscher, R. T. & Knapp, A. N. Global Patterns of Surface Ocean Dissolved Organic Matter Stoichiometry. Glob. Biogeochem. Cycles 37, e2023GB007788 (2023).

27. Fakhraee, M., Tarhan, L. G., Planavsky, N. J. & Reinhard, C. T. A largely invariant marine dissolved organic carbon reservoir across Earth’s history. Proc. Natl. Acad. Sci. U. S. A. 118, e2103511118 (2021).

28. Claire, M. W. et al. The Evolution of Solar Flux from 0.1 nm to 160 μm: Quantitative Estimates for Planetary Studies. Astrophys. J. 757, 95 (2012).

29. Farr, O. et al. Archean phosphorus recycling facilitated by ultraviolet radiation. Proc. Natl. Acad. Sci. 120, e2307524120 (2023).

30. Mayer, L. M., Schick, L. L., Bianchi, T. S. & Wysocki, L. A. Photochemical changes in chemical markers of sedimentary organic matter source and age. Mar. Chem. 113, 123–128 (2009).

31. Kieber, R. J., Whitehead, R. F. & Skrabal, S. A. Photochemical production of dissolved organic carbon from resuspended sediments. Limnol. Oceanogr. 51, 2187–2195 (2006).

32. Ward, C. P., Nalven, S. G., Crump, B. C., Kling, G. W. & Cory, R. M. Photochemical alteration of organic carbon draining permafrost soils shifts microbial metabolic pathways and stimulates respiration. Nat. Commun. 8, 772 (2017).

33. Galili, N. et al. The geologic history of marine dissolved organic carbon from iron oxides. Nature 644, 945–951 (2025).

34. Crockford, P. W., Bar On, Y. M., Ward, L. M., Milo, R. & Halevy, I. The geologic history of primary productivity. Curr. Biol. 33, 4741–4750.e5 (2023).

35. Gregory, B. S., Claire, M. W. & Rugheimer, S. Photochemical modelling of atmospheric oxygen levels confirms two stable states. Earth Planet. Sci. Lett. 561, 116818 (2021).

36. Liu, J., Hardisty, D. S., Kasting, J. F., Fakhraee, M. & Planavsky, N. J. Evolution of the iodine cycle and the late stabilization of the Earth’s ozone layer. Proc. Natl. Acad. Sci. 122, e2412898121 (2025).

37. Pontrelli, S. et al. Metabolic cross-feeding structures the assembly of polysaccharide degrading communities. Sci. Adv. 8, eabk3076 (2022).

38. Ebrahimi, A., Schwartzman, J. & Cordero, O. X. Cooperation and spatial self-organization determine rate and efficiency of particulate organic matter degradation in marine bacteria. Proc. Natl. Acad. Sci. 116, 23309–23316 (2019).

39. Hoppe, H.-G. Phosphatase activity in the sea. Hydrobiologia 493, 187–200 (2003).

40. Srivastava, A. et al. Enzyme promiscuity in natural environments: alkaline phosphatase in the ocean. ISME J. 15, 3375–3383 (2021).

41. Luo, H., Benner, R., Long, R. A. & Hu, J. Subcellular localization of marine bacterial alkaline phosphatases. Proc. Natl. Acad. Sci. 106, 21219–21223 (2009).

42. Reinhard, C. T. et al. Evolution of the global phosphorus cycle. Nature 541, 386–389 (2017).

43. Boden, J. S., Zhong, J., Anderson, R. E. & Stüeken, E. E. Timing the evolution of phosphorus-cycling enzymes through geological time using phylogenomics. Nat. Commun. 15, 3703 (2024).

44. Parsons, C., Stüeken, E. E., Rosen, C. J., Mateos, K. & Anderson, R. E. Radiation of nitrogen‐metabolizing enzymes across the tree of life tracks environmental transitions in Earth history. Geobiology 19, 18–34 (2021).

45. Davín, A. A. et al. A geological timescale for bacterial evolution and oxygen adaptation. Science 388, eadp1853 (2025).

46. Chen, X. et al. Transient marine bottom water oxygenation on continental shelves by 2.65 billion years ago. Nat. Geosci. 18, 423–429 (2025).

47. Liang, X., Stüeken, E. E., Alessi, D. S., Konhauser, K. O. & Li, L. A seawater oxygen oscillation recorded by iron formations prior to the Great Oxidation Event. Nat. Geosci. 18, 417–422 (2025).

48. Czaja, A. D. et al. Iron and carbon isotope evidence for ecosystem and environmental diversity in the ∼2.7 to 2.5Ga Hamersley Province, Western Australia. Earth Planet. Sci. Lett. 292, 170–180 (2010).

49. Konhauser, K. O., Newman, D. K. & Kappler, A. The potential significance of microbial Fe(III) reduction during deposition of Precambrian banded iron formations. Geobiology 3, 167–177 (2005).

50. Konhauser, K. O. et al. Iron formations: A global record of Neoarchaean to Palaeoproterozoic environmental history. Earth-Sci. Rev. 172, 140–177 (2017).

51. Lyons, T. W. et al. Co‐evolution of early Earth environments and microbial life. Nat. Rev. Microbiol. 22, 572–586 (2024).

52. Kipp, M. A. & Stüeken, E. E. Biomass recycling and Earth’s early phosphorus cycle. Sci. Adv. 3, eaao4795 (2017).

53. Fakhraee, M., Planavsky, N. J. & Reinhard, C. T. The role of environmental factors in the long-term evolution of the marine biological pump. Nat. Geosci. 13, 812–816 (2020).

54. Kipp, M. A., Krissansen‐Totton, J. & Catling, D. C. High Organic Burial Efficiency Is Required to Explain Mass Balance in Earth’s Early Carbon Cycle. Glob. Biogeochem. Cycles 35, 2020GB006707 (2021).

55. Lotem, N. et al. Reconciling Archean organic-rich mudrocks with low primary productivity before the Great Oxygenation Event. Proc. Natl. Acad. Sci. 122, e2417673121 (2025).

56. Hao, J. et al. Cycling phosphorus on the Archean Earth: Part II. Phosphorus limitation on primary production in Archean ecosystems. Geochim. Cosmochim. Acta 280, 360–377 (2020).

57. Fakhraee, M., Tarhan, L. G., Reinhard, C. T. & Planavsky, N. J. Constraining the elemental stoichiometry of early marine life. Geology 51, 1043–1047 (2023).

58. Planavsky, N. J. et al. On carbon burial and net primary production through Earth’s history. Am. J. Sci. 322, 413–460 (2022).

59. Lyons, T. W., Diamond, C. W., Planavsky, N. J., Reinhard, C. T. & Li, C. Oxygenation, Life, and the Planetary System during Earth’s Middle History: An Overview. Astrobiology 21, 906–923 (2021).

