## Supplementary material for "Microbial exoenzymes catalyzed the transition to an oxygenated Earth": Methods & Supplemental Information

**Materials & Methods**

**Compiling genomes from distinct microbial metabolic guilds**

We assembled a dataset of high-quality microbial genomes to assess the distribution of exoenzymes across distinct metabolic guilds: aerobic heterotrophy, fermentation, dissimilatory iron reduction, dissimilatory sulfate reduction, dissimilatory nitrate reduction, methanogenesis, anoxygenic photoautotrophy, and oxygenic photoautotrophy. Genomes were retrieved from the Joint Genome Institute’s (JGI) genomic catalog of Earth’s microbiomes (GEMs) and the National Center for Biotechnology Information (NCBI) database based on the presence of one or more marker genes that correspond to each microbial metabolism (**Table S1**). In the case of oxygenic photoautotrophy, we used genomes that were classified as Cyanobacteria based on taxonomy via 16S rRNA sequences. Fermenters do not possess well-established marker genes. We retrieved genomes belonging to fermenters using a dataset compiled by Hackmann and Zhang (2023) that includes phenotypic and genotypic records of 4355 prokaryotic fermentative organisms. For aerobic heterotrophy, we focused our efforts on obtaining genomes that reflected modern marine environments. We retrieved genomes from two sources – a culture collection of marine bacteria isolated from coastal seawater (Gralka et al. 2023), and MAGs from the GEMs catalog and Tara Oceans dataset (Tully et al. 2018). To identify MAGs of aerobic heterotrophs, we confirmed the absence of marker genes specific to autotrophy and verified the presence of cytochrome *c* oxidase, an enzyme that reduces oxygen in the final step of the electron transport chain and is a key component of aerobic respiration.

To detect marker genes in genomes from the NCBI and GEM databases we used well-established hidden Markov model (HMM) profiles of the relevant molecular marker genes (**Table S2**). These HMM profiles were obtained from the NCBI’s protein family model pages or were published in the literature (Garber et al. 2020, Appler et al. 2025). We applied e-value thresholds above the full-sequence score for each HMM. For the *coxABC* HMM, where a sequence score was not available, an e-value threshold of 1x10^-50^ was used (Appler, personal communication). Marker genes in genomes were identified using the *hmmsearch* command from HMMER 3.3.2 (Eddy, 2011). Genomes containing marker genes were processed using a custom R script, rtackhmmer which is available on GitHub. The completeness and contamination of these genomes were assessed with CheckM v1.1.6 (Parks et al. 2015). Only genomes over 90% complete and with less than 1.3% contamination were retained, exceeding the high-quality level of the minimum information about a metagenome-assembled genome (MIMAG) standard (Bowers et al. 2027). Where possible, we selected genomes isolated from aquatic and terrestrial environments while excluding those from engineered environments (e.g. fracking water, fracture fluid, mine pit ponds, drinking water treatment plant) based on the provided metadata. Lastly, we re-annotated all the compiled genomes using Prokka 1.14.6 (Seemann, 2014) with a stringent e-value threshold of 1x10^-50^ to maintain a conservative and uniform annotation standard.

*Identification of exoenzymes in genomes*

To identify putative exoenzymes in the compiled genomes, we searched for 22 terms that corresponded to well-known hydrolytic enzymes (**Table S3**) using Geneious (v2023.2.1). The resulting gene sequences of identified enzymes were translated into protein sequences. All domains of life use the secretion (Sec) and twin arginine translocation (Tat) pathways to export enzymes across cellular membranes (**Fig. S3**; Robinson & Bolhius, 2004; Papanikou et al. 2007). Diverse methods of protein secretion have been identified in microbes, including those that work in conjunction with the Sec and Tat pathways (Type II & V), and those that export proteins from the cytoplasm into the environment (Type I) or recipient cells (Type III, IV & VI) directly (Green & Mescas, 2016). We focused our efforts on exoenzymes secreted through the Sec and Tat pathways, which are likely the oldest (Moody et al. 2024; Wu et al. 2000) and most ubiquitous (Pugsley et al. 2004) transport systems. The ability of microorganisms to export enzymes using the Sec and Tat pathways is governed by a short stretch of amino acids referred to as the signal peptide sequence, which allows putative exoenzymes to be identified from their amino acid sequence (Owji et al. 2018). The presence of signal peptide sequences was detected using SignalP 6.0 (Teufel et al. 2022). However, membrane proteins and proteins residing in the periplasmic space also require a signal peptide to cross the inner membrane. To determine whether signal-peptide containing sequences were likely to encode a secreted enzyme, we predicted protein localization using PSORTb v3.0 (Yu et al. 2010). PSORTb is a generalist protein localization software, employing multiple strategies to reach a prediction. Importantly for our purpose, it focuses on high precision to minimize false positives at the expense of assigning predictions to all sequences. Prior to processing protein sequences through PSORTb, sequences were separated by predicted cell type (Gram stain) based on taxonomy. Taxonomy predictions were taken from NCBI and GEM metadata. For phyla without cultured representatives, cell type was taken from Witwinowski et al. (2022), or, if unclear or unknown based on the literature, assumed to be gram negative. Thus, only enzyme sequences that were both predicted to contain a signal peptide and identified as extracellular based on their predicted structure were included in the final dataset. All predicted exoenzymes were subject to manual inspection to exclude those involved in housekeeping roles such as maintaining cellular membrane.  A full list of removed and included protein sequences is provided (**Tables S4**). The workflow for compiling genomes and predicting exoenzymes is summarized in **Figure S4**.

**Clustering of exoenzymes sequences using sequence similarity networks**

To further explore the evolutionary and functional relationships between the predicted exoenzymes, sequence similarity networks (SSNs) were generated for (1) all of the predicted exoenzyme complete sequences (including their signal peptide). In SSNs, each sequence is represented as a single point (“node”) and connected to other nodes through “edges”, according to a chosen threshold e-value. Groups of highly similar sequences group together in clusters thereby mimicking clade formation as seen in traditional phylogenetic trees.  To generate the SSNs, a pairwise similarity comparison was performed separately for (1) all predicted exoenzyme and (2) signal peptide sequences (Blast all-against-all; Altschul et al. 1997). Each pairwise comparison generates an e-value, which are then used to construct a network in Cytoscape (Shannon et al. 2003). An edge was drawn between two nodes if the e-value is equal or lower than 1x10^-30^. The length of connecting edges correlates to the relative dissimilarity of a given sequence pair, whereas the relative positioning of clusters holds no meaning. Thresholds for e-values were determined by testing various cutoff values for connecting edges (10^-10^, 10^-15^, 10^-20^, 10^-25^, and 10^-30^) and visually assessing the balance between defined similarity-based clusters and the representation of associations with the applied variables. The networks were visualized in Cytoscape using the organic layout, with enzyme type and metabolic guild of the source organism (Teufel et al. 2022), mapped onto both networks as node color.

**Ancient Ocean Carbon Model**

The modeling framework used in this study builds upon a previously published global ocean box model that simulates the production, transport, and degradation of dissolved and particulate organic carbon through geological time (Fakhraee et al. 2021). This earlier model coupled a simplified ocean-atmosphere carbon cycle to biological and physical processes regulating organic matter cycling and was used to investigate the evolution of marine DOC in the Precambrian. The model divides the ocean into a series of interconnected boxes representing surface and deep layers across different oceanic regions (e.g. open ocean, coastal shelf, marginal seas), with mass-balance equations governing the cycling of carbon between these compartments. DOC and POC reservoirs in each box are dynamically updated through inputs from primary production, outputs via remineralization and burial, and transport across boxes through advection and diffusion.

The biological pump is explicitly modeled via a stochastic particle aggregation scheme, where particles collide, stick, and sink according to size-dependent velocities. POC degradation during transit is described using a power-law remineralization profile with parameters tuned to reflect particle flux attenuation under different redox and thermal conditions. DOC remineralization is modeled as a Monod-type uptake process, where microbial consumption of DOC is proportional to substrate concentration and limited by microbial processing capacity. Partitioning between labile and recalcitrant DOC pools is included, allowing for differential turnover times. The model also incorporates riverine, hydrothermal, and sedimentary sources of DOC, and accounts for abiotic production pathways such as photodegradation of POC in the surface ocean.

In this study, we expanded the model to explore how UV-driven photochemical processes might have influenced DOC dynamics under early Earth conditions. Specifically, we incorporated a refined parameterization of UV-dependent DOC production based on enhanced solar UV flux in the absence of an ozone layer. To do so, we simulated the UV flux available for photodegradation in the surface layer of Earth's ocean throughout geological history. The Sun's irradiance spectrum throughout geological time, as described by Claire et al. (2012) was interpolated using a univariate spline for three different eons (**Fig. S7**). The eons were defined as follows: Archean (4 Ga to 2.4 Ga), Proterozoic (2.4 Ga to 540 Ma) and Phanerozoic (540 Ma to present day).

Next, we modelled the irradiance spectrum at Earth’s surface over time to account for changes in Earth’s atmosphere that directly influence UV flux reaching the surface. We assumed that the total UV flux at the top of the atmosphere increases monotonically over time and adopted a time resolution of 0.05 giga years (50 million years), and the same wavelength resolution as Claire et al. (2012). The atmosphere was modelled considering only carbon dioxide (CO_2_), nitrogen (N_2_), oxygen (O_2_), and ozone (O_3_) (Cnossen et al. 2007). For the Archean eon, we assumed a total pressure of 0.23 bar with 70% CO_2_ and 30% N_2_ (Som et al. 2016; Lehmer et al. 2020). The total atmospheric pressure and abundances of CO_2_ and N_2_ were assumed to change linearly from 2.7 Ga values (Lehmer et al. 2020) to modern values of 1.01 bar, 0.04% CO_2_, and 78% N_2_. This allowed us to estimate the atmospheric pressure and abundances of CO_2_ and N_2_ for the Proterozoic and Phanerozoic. O_2_ concentrations over time were obtained from Lyons et al. (2014). The thickness of the ozone layer over time was calculated using O_2_ concentrations (Lyons et al. 2014) and their relationship to the O_3_ column (Cooke et al. 2022).

Combining the time-evolving atmospheric abundance (**Fig. S8**) and the appropriate absorption cross-sections from Cnossen et al. (2007), we calculated the atmosphere’s optical depth spectrum (**Fig. S9**) and the irradiance spectrum at Earth’s surface over time (**Fig. S10**). Then, we modelled UV light propagation in the ocean using attenuation coefficients for water (Hale & Querry, 1973; **Fig. S11**) and subsequently applied those values to calculate the irradiance spectrum in the ocean's mixed layer at the three eons (**Fig. S12**). We assumed that ocean water optical properties remain constant throughout Earth's history. Lastly, to assess the robustness of our approach, we examined how atmospheric composition during different eons, and the depth of the ocean's mixed layer, affected our results. Using different models of O_2_ or CO_2_ evolution in the atmosphere did not qualitatively affect the irradiance spectra in the ocean’s mixed layer. Moreover, changing the depth of the ocean's mixed layer (varied between 10 m to 50 m), the total atmospheric pressure (from 0.23 bar and 1.20 bar), and the proportion of CO_2_ to N_2_ (from 70:30 to 30:70), did not qualitatively change the irradiance spectra in the ocean’s mixed layer.

Photochemical transformation rates were scaled with estimates of Archean UV irradiance, and we included constraints on POC availability arising from lower net primary productivity in the early biosphere (100–1,000× lower than today). These updates allowed us to examine the interplay between DOC supply, microbial abundance, and enzymatic demand under Precambrian environmental conditions. Model outputs were validated against modern DOC and POC distributions and evaluated against observations from low-productivity and redox-stratified analog environments such as the Black Sea and Lake Pavin. This framework allowed us to explore how the ratio of DOC to POC and DOC to microbial cell abundance evolved over Earth history, and what implications these patterns hold for the development of microbial hydrolytic strategies and carbon turnover in the ocean.

**Alkaline phosphatase molecular clock**

Our tree of life (“species tree”) was constructed previously (Boden et al. 2024) using sixteen ribosomal proteins fromf 865 genomes chosen to represent the breadth of Earth’s microbial diversity. Relaxed Bayesian molecular clocks were implemented in Phylobayes v.4.1 (Lartillot et al. 2009) to anchor this species tree to a geological timeline using eight calibration points based on the fossil record (Boden et al., 2024). To reflect different model assumptions regarding the inheritance of substitution rates between lineages, we estimated divergence times using two autocorrelated clock models (lognormal (LN) and cox-ingersoll-ross (CIR)) and the uncorrelated gamma multipliers (UGAM) clock model. A full description of the parameters is presented in Boden et al. 2024, but here we applied an earlier hard minimum calibration for the origin of methanogenesis at more than 3.46 Ga based on new geochemical and biological data (Cavalazzi et al. 2021; Ueno et al. 2006; Wolfe & Fournier, 2018). Two independent chains were implemented for 129,168 to 273,674 cycles for each molecular clock model until a pair of chains reached convergence, defined as relative differences of all parameters < 0.3 and effective sizes > 50 after the first 25% of cycles had been discarded as burn-in. These relative differences and effective sizes were calculated with tracecomp implemented in Phylobayes v.4.1 (Lartillot et al. 2009).

We used HMMER (Eddy, 2011) to search for homologs of alkaline phosphatase genes *phoX, phoA,* and *phoD* within the genomes we used for the species tree. HMM profiles obtained from the Pfam database and NCBI’s protein family model database (**Table S5**) and an e-value threshold of 0.001 were applied. Of these three alkaline phosphatase enzymes, only the *phoD* protein tree had monophyletic clades of extracellular sequences (**Fig. S1**). As such, only the methods for *phoD* are described below. Non-monophyletic clades comprise a mixture of sequences with varied subcellular localizations alkaline phosphatase sequences such that some gene events may correspond to **intracellular, rather than extracellular enzyme function.** HMMER3 hits for each enzyme were aligned with the query sequences using MAFFT v.7.4 (Katoh & Standley, 2013) with the high accuracy progressive alignment strategy (E-large-INS-1, Nakamura et al. 2018) designed for alignments containing multiple regions of alignable residues separated by unalignable residues and gaps (command --mpi --large --genafpair). Phylogenetically uninformative positions defined as containing more than or equal to 85 % gaps were removed using trimal v.1.4.rev22 (Capella-Gutierrez et al. 2009). The resulting trimmed alignments were then used to reconstruct a rooted maximum likelihood tree detailing the relationships between *phoD* and its homologs using IQTREE v.2.2.5 (Minh et al. 2020) with the best of four non-reversible complex substitution models (namely NQ.pfam+C20+G4, NW.pfam+C20+G4+F, NQ.bac+C20+G4,NQ.bac+C20+G4+F) chosen by ModelFinder (Kalyaanamoorthy et al. 2017). Branch supports were estimated by calculating ultrafast bootstraps with 1000 replicates (Hoang et al. 2017) and up to 5000 iterations (-bb 1000 -nm 5000). Tree topology tests, including the approximately-unbiased (AU) test (Shimodaira, 2002) were applied to measure confidence in the root placements (--root-test -zb 1000 -au). The tree was rooted on the branch with the highest bootstrap support values (77.8) and which passed the AU test (defined as AU p-values more than or equal to 0.1). Any homologs which did not descend from most recent common ancestor of known enzymes were removed. In this case, 227 homologs of PhoD were retained for further analyses, and 1 was removed. The remaining 227 homologs were then re-aligned using a more accurate iterative alignment method (namely E-INS-I in MAFFT v7.4 (Katoh & Standley, 2013)), poorly sequenced regions removed with TAPER v.1.0.2 (Zhang et al., 2021), and trimmed less stringently (alignment columns with ≥ 95% gaps were removed with trimal v.1.4.rev22 (Capella-Gutierrez et al., 2009) to retain as much phylogenetic signal as possible. Armed with this more accurate data, a new tree was reconstructed in IQ-TREE v2.0.3 (Minh et al., 2020) with 1,000 ultrafast bootstraps (Hoang et al., 2017) and a broader set of reversible complex mixture models (including LG+G4+C20+F , LG+G4+C60+F, LG+R4+C20+F, LG+R4+C60+F, Q.pfam+G4+C20+F, Q.pfam+G4+C60+F, Q.pfam+R4+C20+F, Q.pfam+R4+C60+F, Q.bac+G4+C20+F, Q.bac+G4+C60+F, Q.bac+R4+C20+F or Q.bac+R4+C60+F). We determined which of the filtered 227 homologs were likely to encode secreted proteins using the same approach as above (section entitled ‘Identification of exoenzymes in genomes’), and annotated them on the *phoD* gene tree. Three monophyletic clades of *phoD* genes were predicted to localize extracellularly, and retained for further analysis while the rest were discarded (**Fig. S1**).

We estimated when these extracellular *phoD* genes spread through microbial lineages by reconstructing three final gene trees, one for each clade of extracellular *phoD* that had been identified. Like before, these reconstructions were made by aligning the genes in MAFFT, removing poorly sequence regions with TAPER and trimming out columns with poor data (defined as ≥ 95 % gaps). Bayesian software (namely MrBayes v. 3.2.7a (Ronquist et al., 2012) with a mixed amino acid model prior, invariant sites and gamma-distributed rates) was used to reconstruct the evolutionary histories instead of IQ-TREE because the sample of trees generated is a better reflection of topological uncertainty. Chains were considered converged when the average standard deviation of split frequencies was < 0.01, the partial scale reduction factor lay between 1.00 and 1.02, and the effective sample size scores of all parameters were > 200 after discarding the first 25% of iterations as burn in. Representative samples of each extracellular *phoD* gene tree were reconciled with the CIR clock model time-calibrated species tree using ecceTERA v. 1.2.5 (Jacox et al. 2016) to identify gene transfer, duplication, loss, and speciation events. The results presented use the default event costs (horizontal gene transfer = 3, duplication = 2, loss = 1, speciation = 0, transfers to the dead allowed) and amalgamated the gene trees (amalgamate = true). The cost associated with a horizontal gene transfer event is based on the resulting impact of transfer, relative to that of duplication, loss, and speciation, on the change in genome size from the parent to the daughter lineages (David and Alm, 2011). The default ratio of 3:2:1:0 would have theoretically differed with variation in genome size, which is unconstrained for early Earth microorganisms. Therefore, to further test how ecceTERA’s reconciliation parameters impact the estimated histories of genes related to alkaline phosphatase and to determine the upper and lower bounds of our estimates, we performed reconciliations with horizontal gene transfer costs of 2, 4, and 6 for all three clock models (CIR, UGAM, and LN) (**Tables S5, S6, S7, S8**). The symmetric median reconciliation from each gene tree was used to determine dates of gene events, confidence intervals, and divergence times using custom Python scripts, which were developed and implemented to time the evolution of sulfur and reduced P species cycling (Mateos et al. 2023; Boden et al. 2024). These scripts were applied using Python v. 2.7.5., ensuring that only reception events were counted for each occurrence of horizontal gene transfers. All trees were visualized in TreeViewer (Bianchini et al. 2024).

**Supplemental Figures**

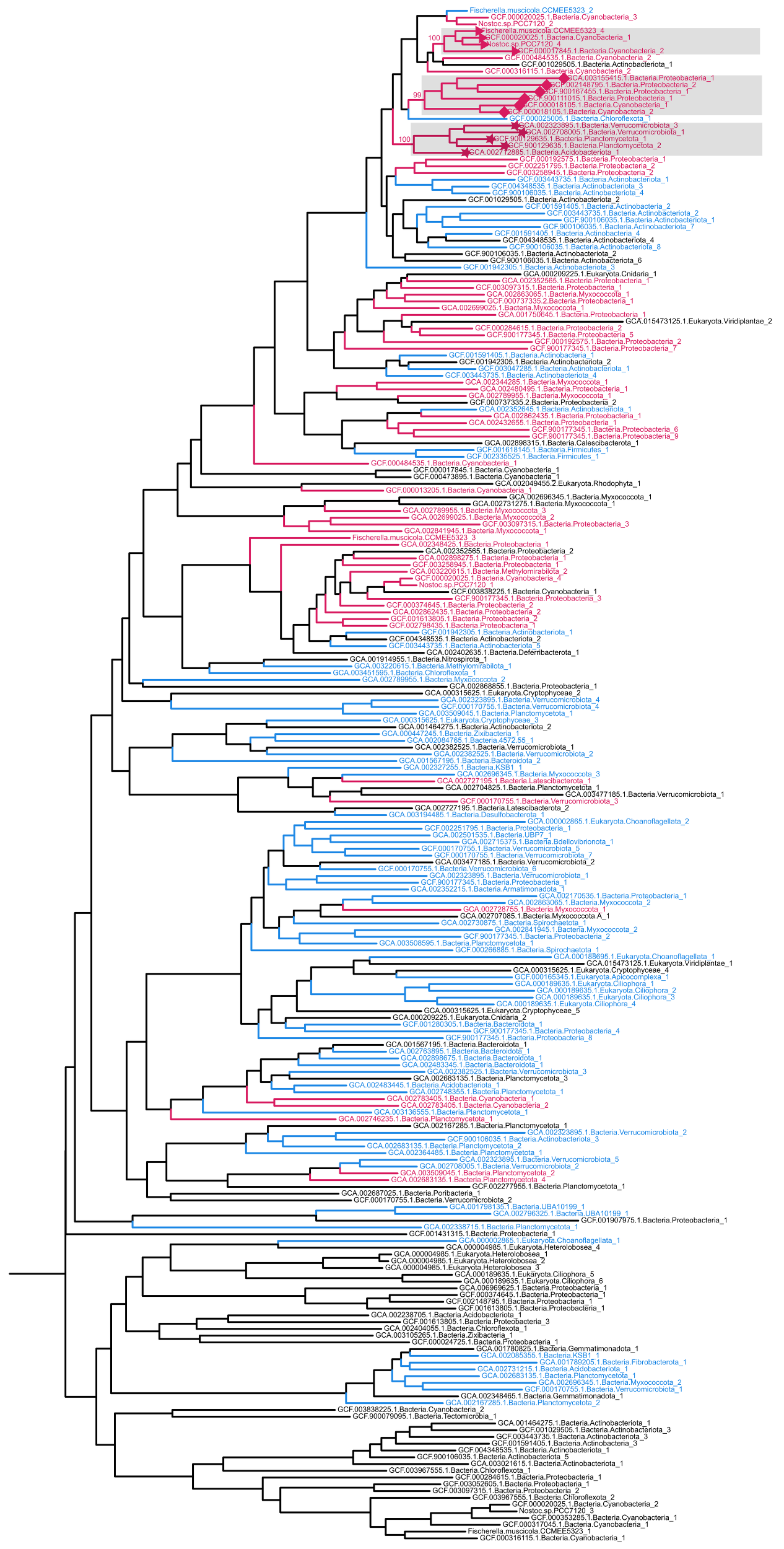

**Figure S1.** Evolutionary tree of PhoD homologs used for reconciliation analyses, with extracellular clades highlighted in grey boxes. Homologs in pink contain predicted signal peptide secretion sequences and are predicted to localize extracellularly. Homologs in blue contain predicted signal peptide secretion sequences but are not predicted to localize extracellularly. Homologs in black do not contain predicted signal peptide secretion sequences.

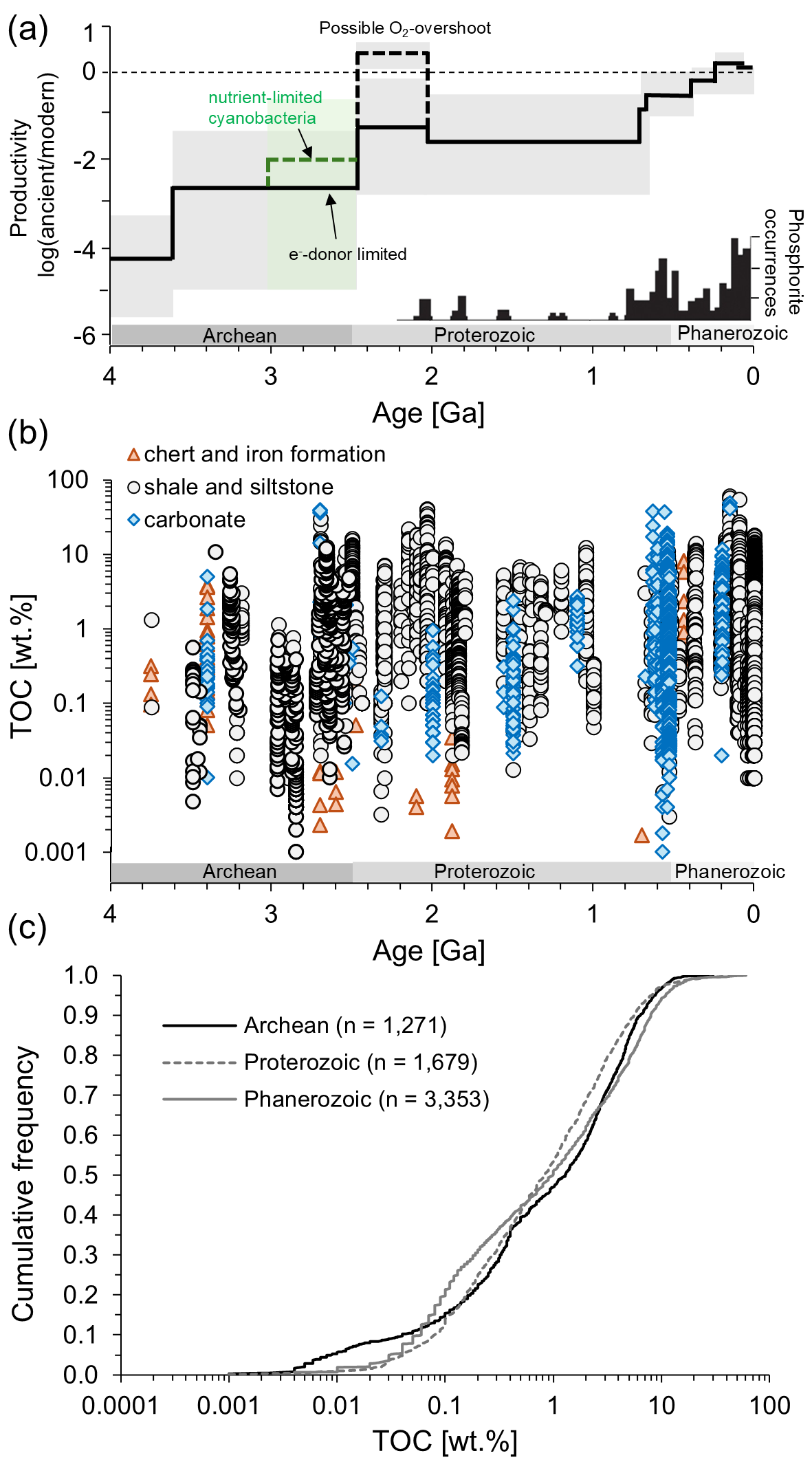

**Figure S2.** (a) Primary productivity through time, redrawn from Crockford et al. (2023). The black line is the most probable value, the shaded regions mark upper and lower limits. (b) Total organic carbon (TOC) in marine sediments through time, compiled from two previous databases (Stüeken et al. 2012; Stüeken et al. 2024). (c) Cumulative frequency distribution of the data shown in panel (b), subdivided into the Archean, Proterozoic and Phanerozoic, showing that the three time periods are effectively indistinguishable from each other in terms of total TOC content.

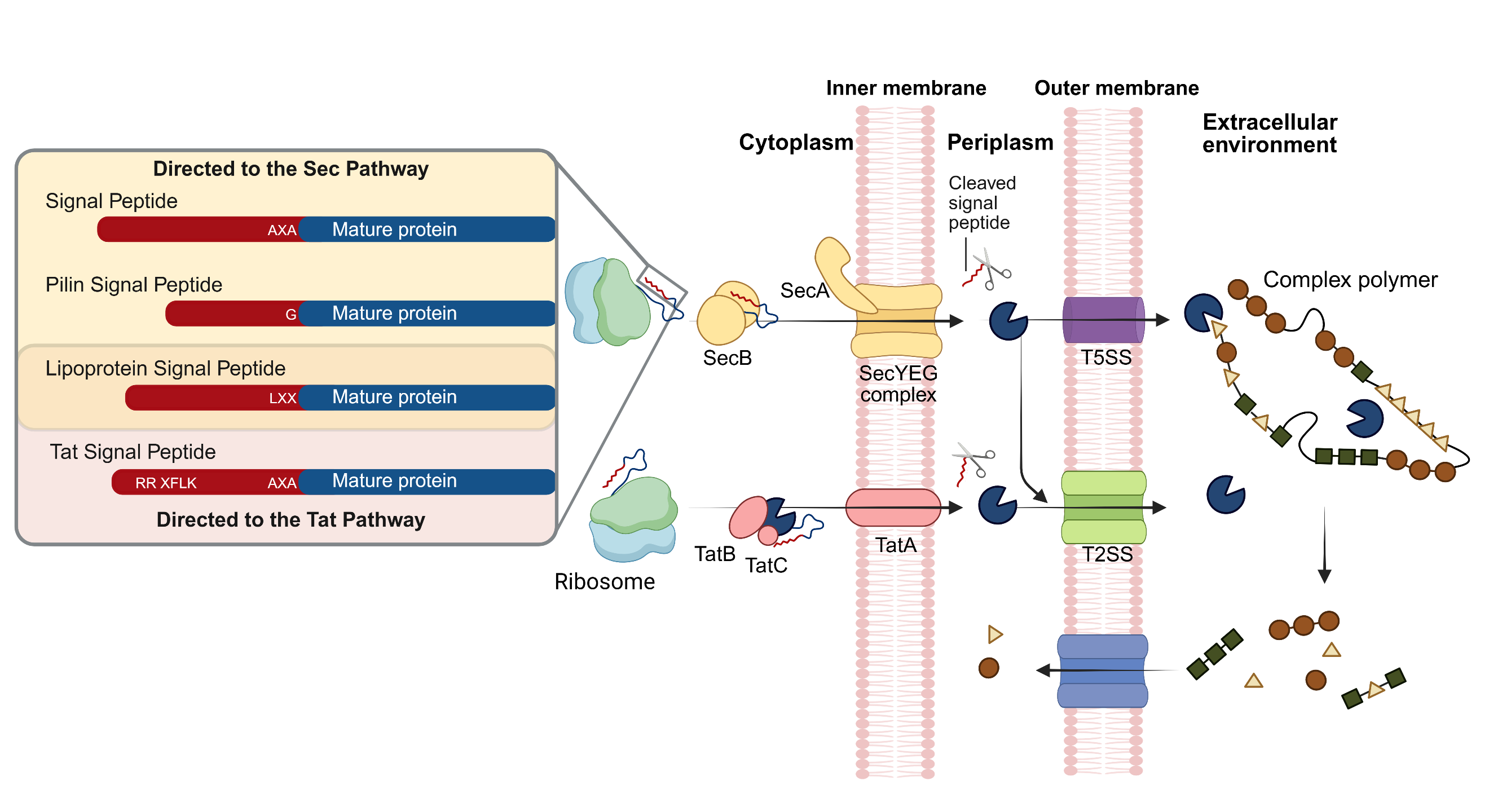

**Figure S3.** **Schematic overview of the transport of secreted proteins in Gram-negative bacteria.** Preproteins are directed to either the Sec (yellow) or Tat (pink) translocation pathways depending on the identity of their N-terminal signal peptide sequence. (left panel). The unfolded Sec-dependant proteins containing a Sec-, prepilin- or lipoprotein-type signal peptide, are translocated by two mechanisms (Almagro Armenteros et al. 2019). The nascent polypeptide chain can be stabilized by SecB after it is released from the ribosome, then targeted to SecA in the inner membrane and translocated by the SecYEG complex across the plasma membrane. Alternatively, SecA can directly interact with the ribosome-bound preprotein and target it to the SecYEG complex for transport. Under both mechanisms, signal peptidase enzymes cleave the signal peptides from the preprotein after translocation. It is only upon transport through the SecYEG complex that the preprotein folds, acquiring tertiary structure. In contrast, proteins directed to the Tat pathway are folded in the cytoplasm and bound by TatB and TatC which then recruit TatA to the inner membrane before translocation. Proteins containing a Tat or lipoprotein signal peptide are targeted to the Tat pathway. Once exported across the inner membrane, (pre)proteins in the periplasm can be transported through Type-2 transporters to the extracellular milieu. Some preproteins that have travelled through the Sec pathway contain the necessary domains to autotransport, folding to form an outer membrane channel that allows passage of the functional part of the protein outside of the cell. Adapted from Kaushik et al. (2022).

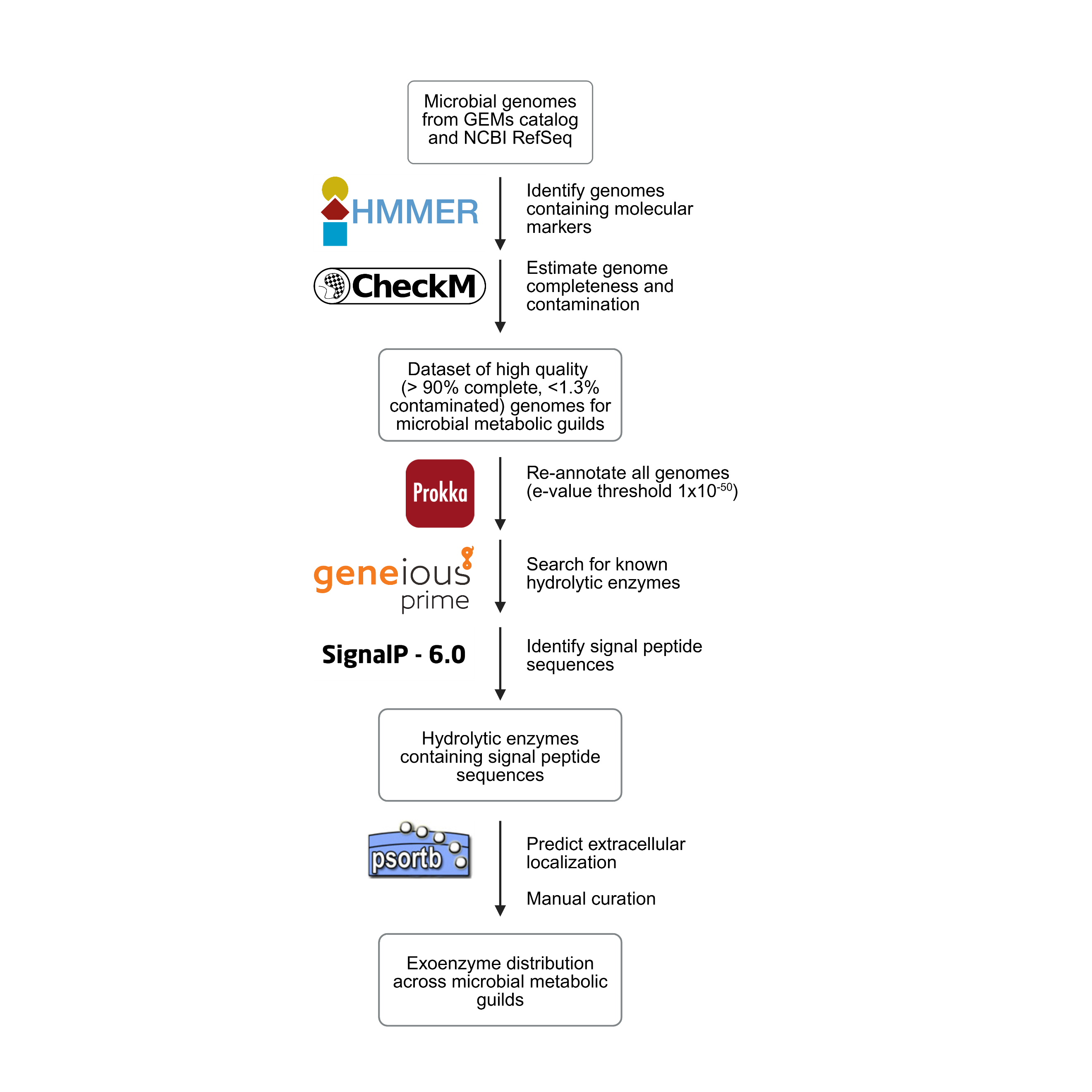

**Figure S4.** Simplified schematic overview of the approach used to predict the presence of exoenzymes in the microbial genomes compiled in this study.

(a)

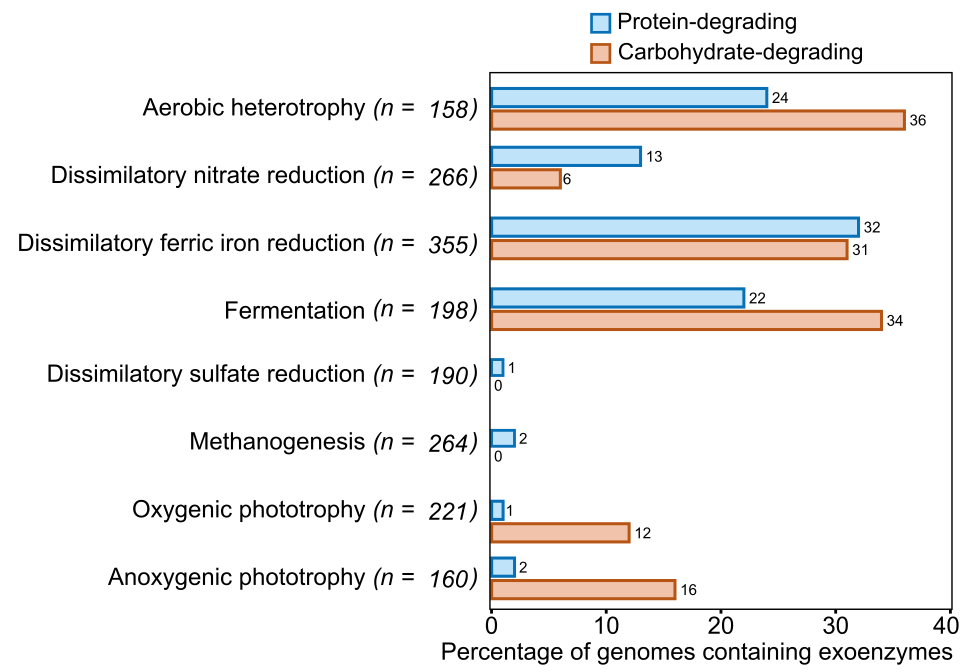

(b)

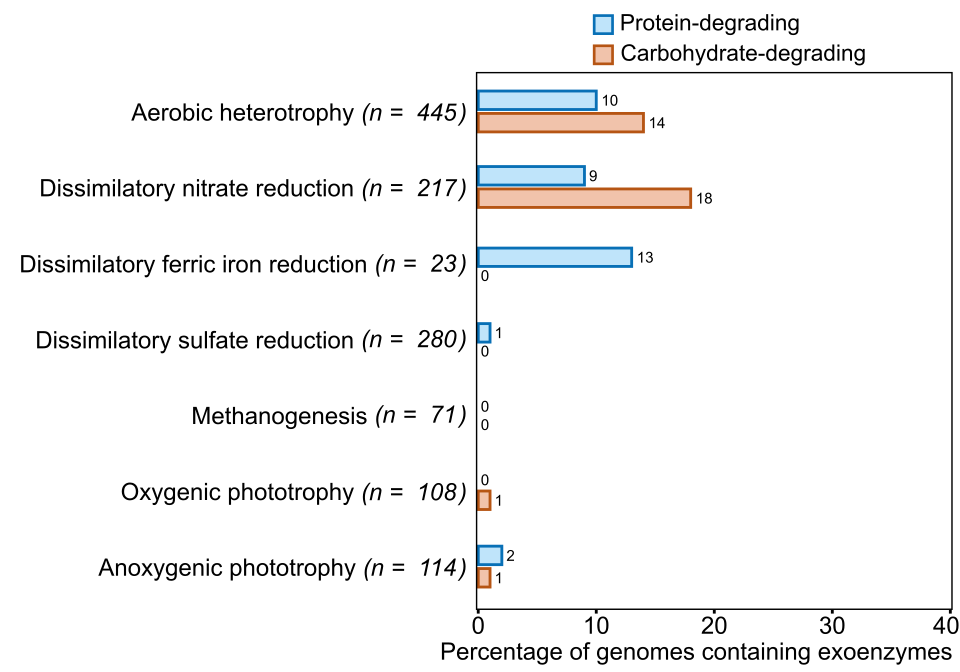

**Figure S5.** The percentage of genomes that contain at least one predicted protein-degrading (blue) or carbohydrate-degrading (orange) exoenzyme in each metabolic group. (A) NCBI RefSeq genomes from metagenome assembled genomes (MAGs) and cultured isolates and (B) JGI GEMs MAGs from aquatic and terrestrial environments.

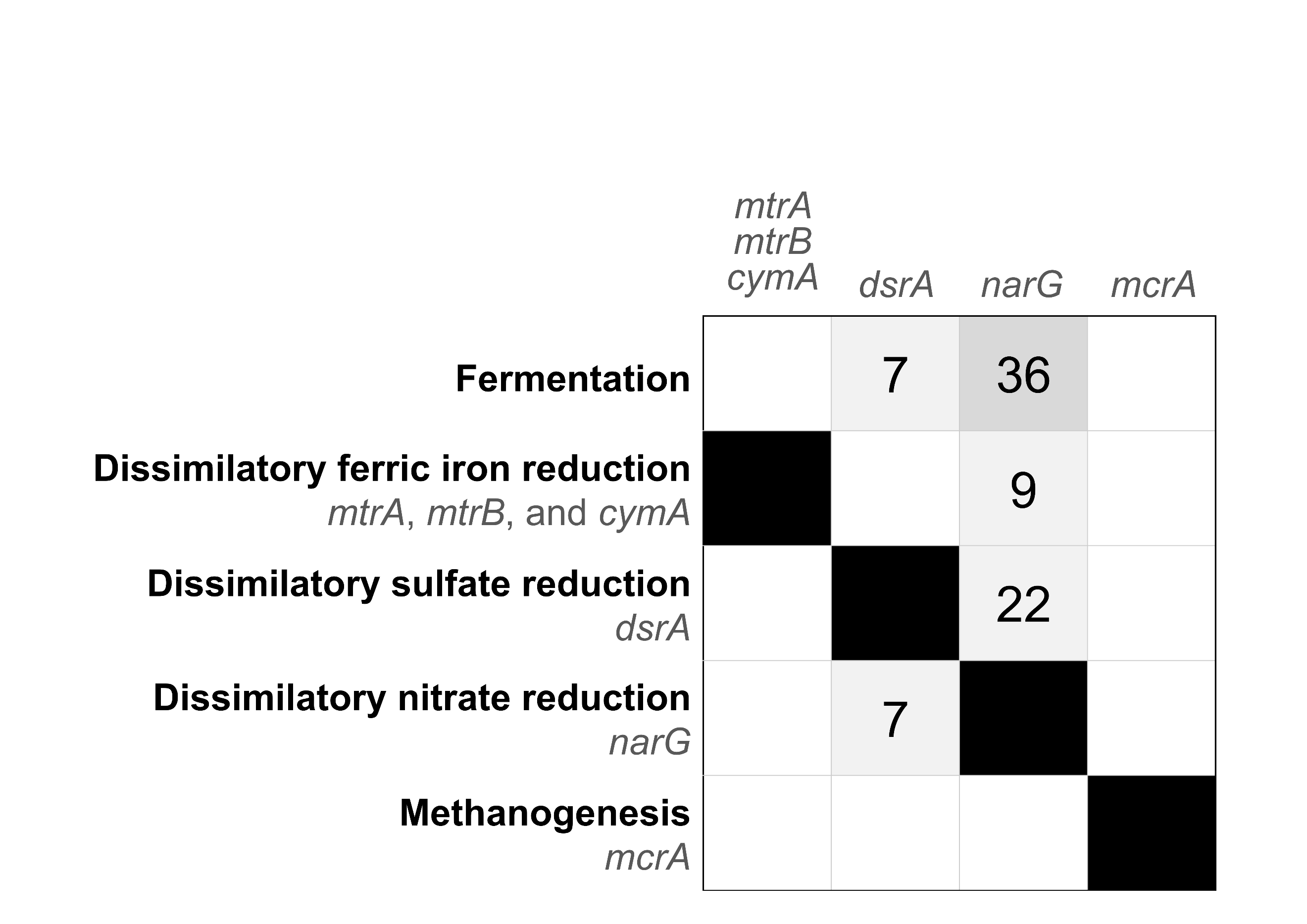

**Figure S6.** Overlap (percentage %) between molecular marker gene(s) in anaerobic heterotrophic genomes as determined using the bit score cutoffs defined in Table S2. Under 5% overlap is omitted for clarity.

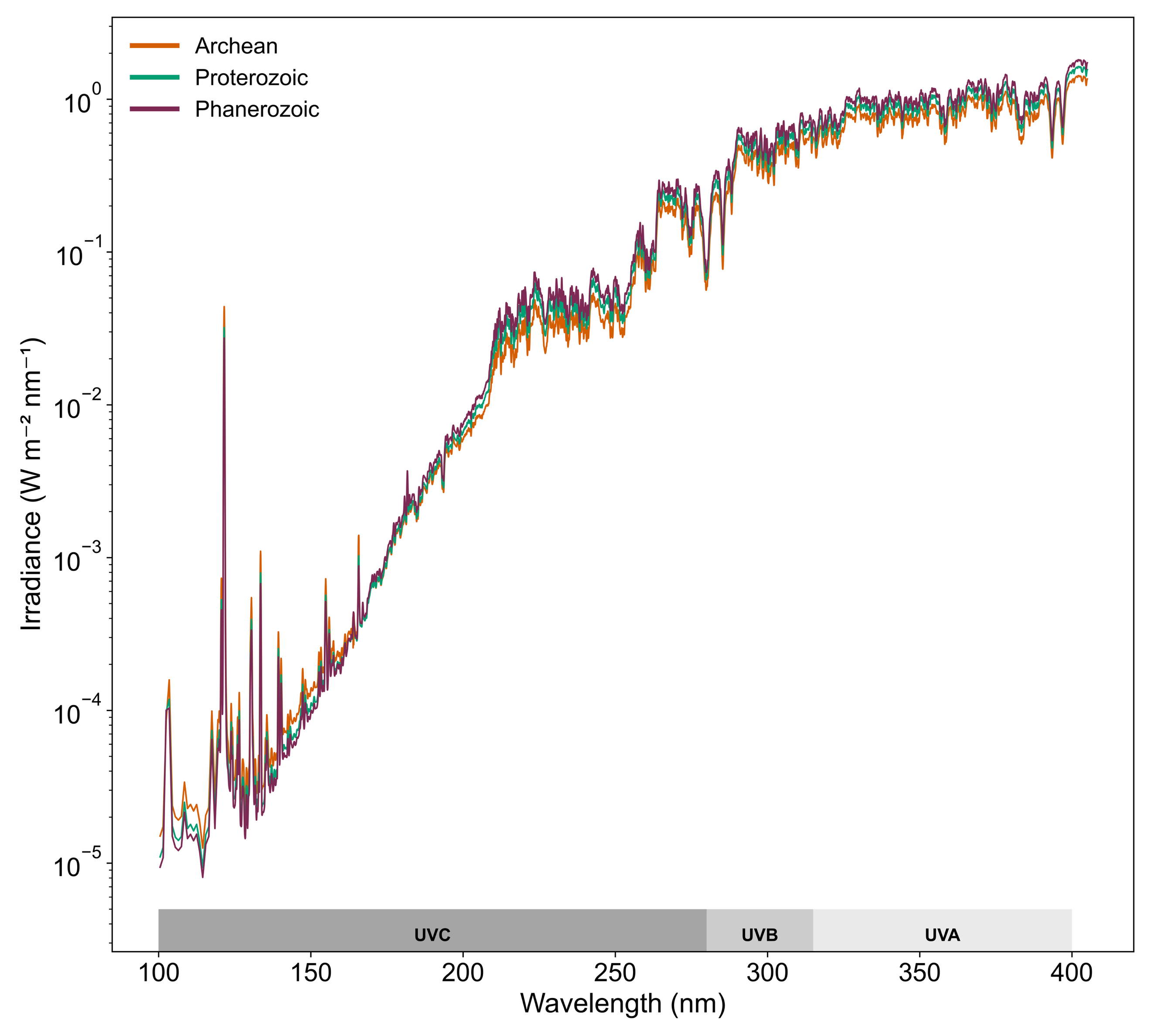

**Figure S7.** Irradiance spectrum of the Sun at the top of Earth’s atmosphere, shown for the Archean, Proterozoic, and Phanerozoic eons by interpolating data from Claire et al. (2012).

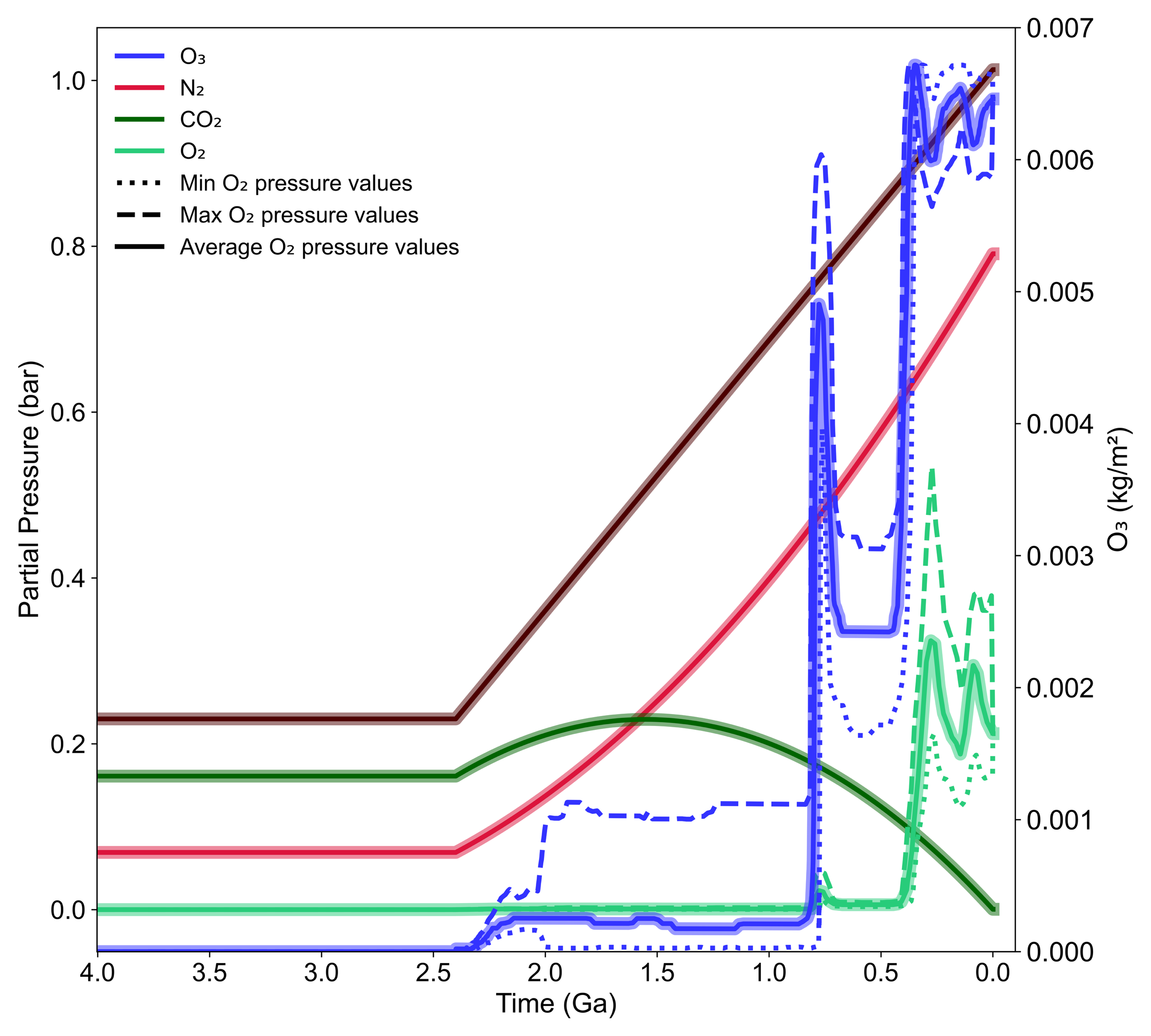

**Figure S8.** Changes in Earth’s atmospheric composition over geological time. CO₂, N₂, and O₂ are expressed as partial pressures, and O₃ is expressed as mass per unit area. O₂ and O₃ values incorporate the upper and lower bounds of O₂ partial pressure from Lyons et al. (2014).

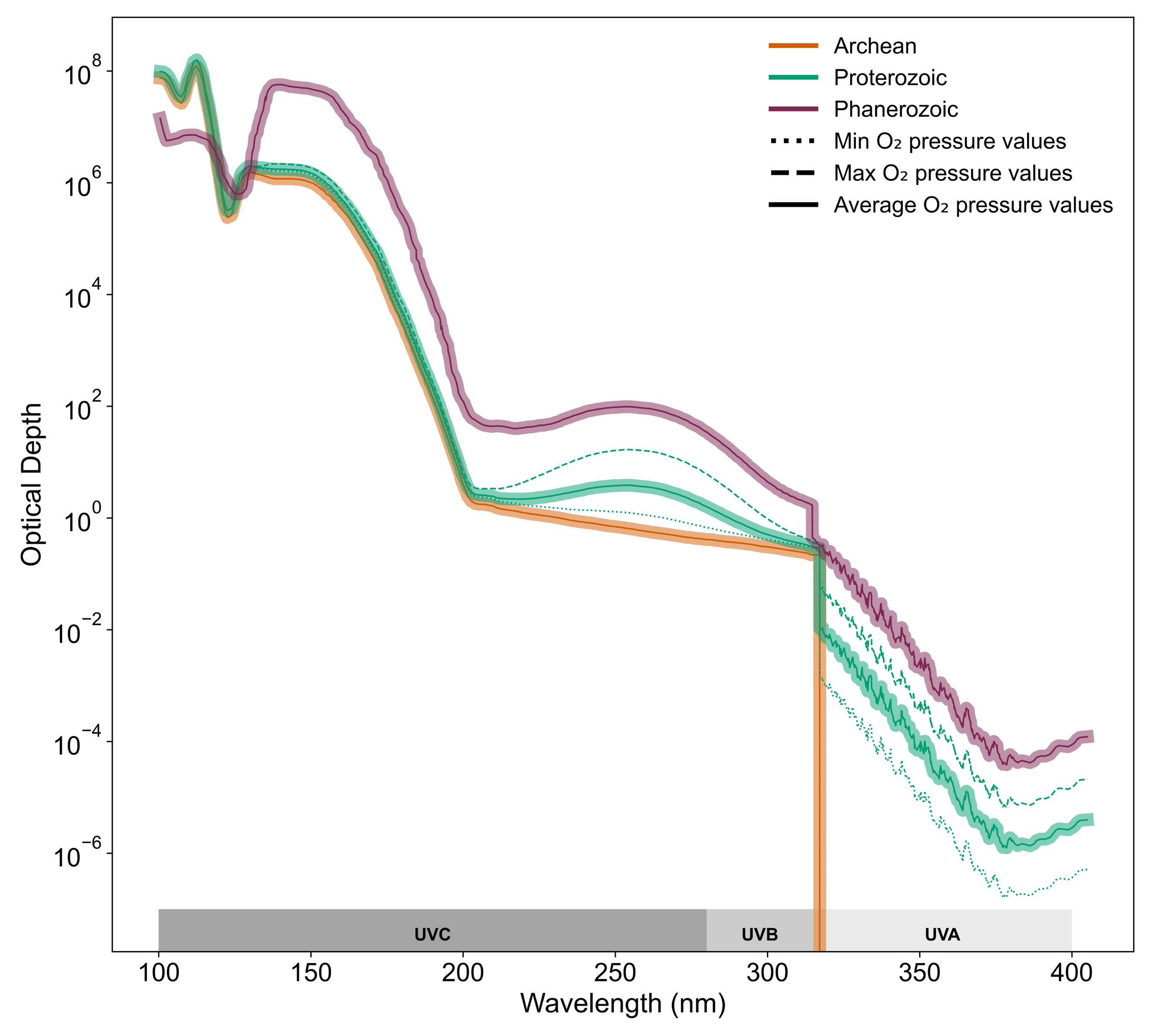

**Figure S9.** Atmospheric absorption spectrum calculated from the modeled atmospheric composition using CO₂, N₂, O₂, and O₃ absorption cross-sections from Cnossen et al. (2007) for the Archean, Proterozoic, and Phanerozoic. Absorption is expressed as optical depth, which describes the atmosphere’s opacity at a given wavelength.

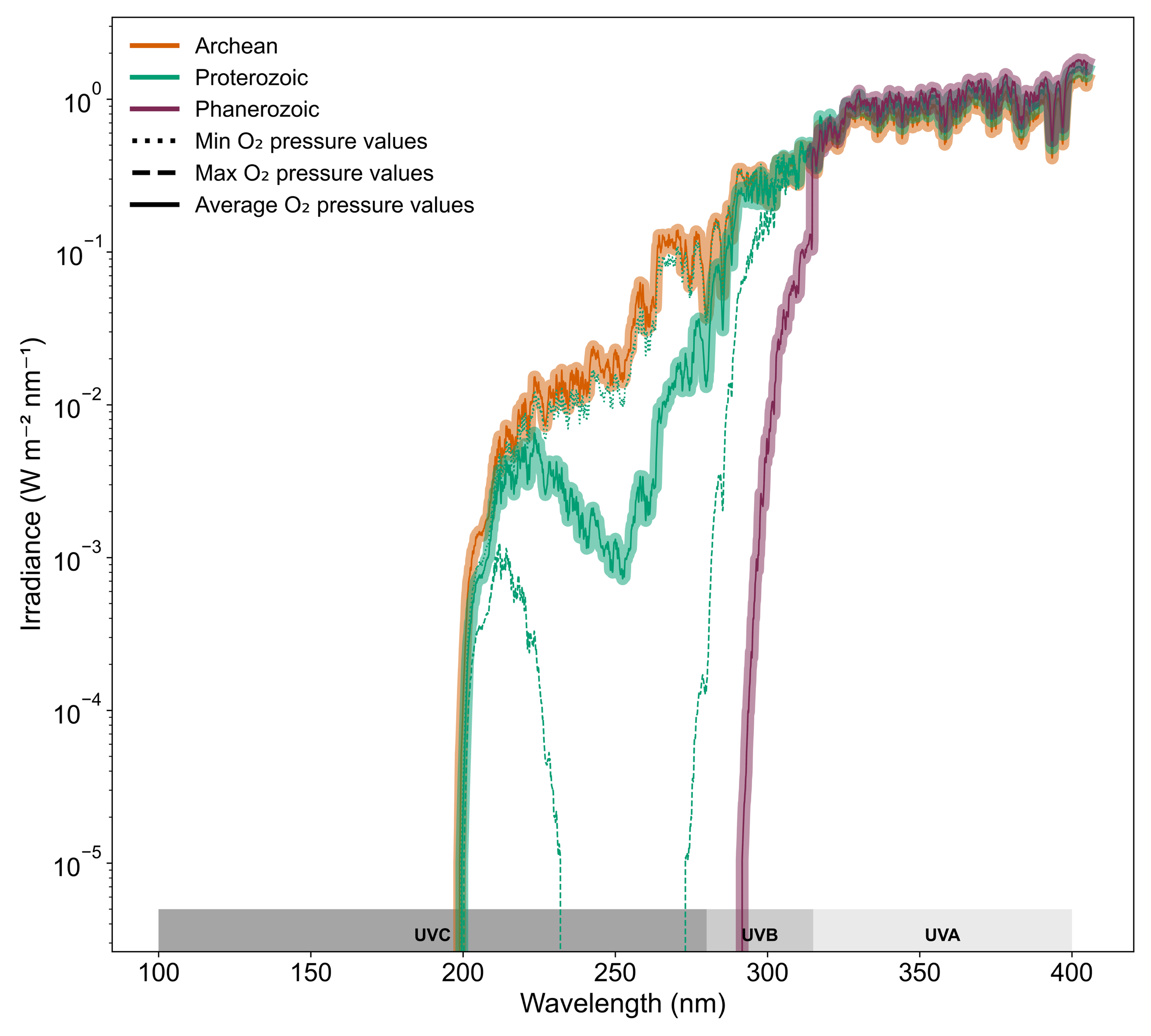

**Figure S10.** Irradiance spectrum at Earth’s surface for the Archean, Proterozoic, and Phanerozoic. The spectra were obtained by applying the atmosphere’s optical depths to the irradiance spectra at the top of Earth's atmosphere using the Lambert-Beer law.

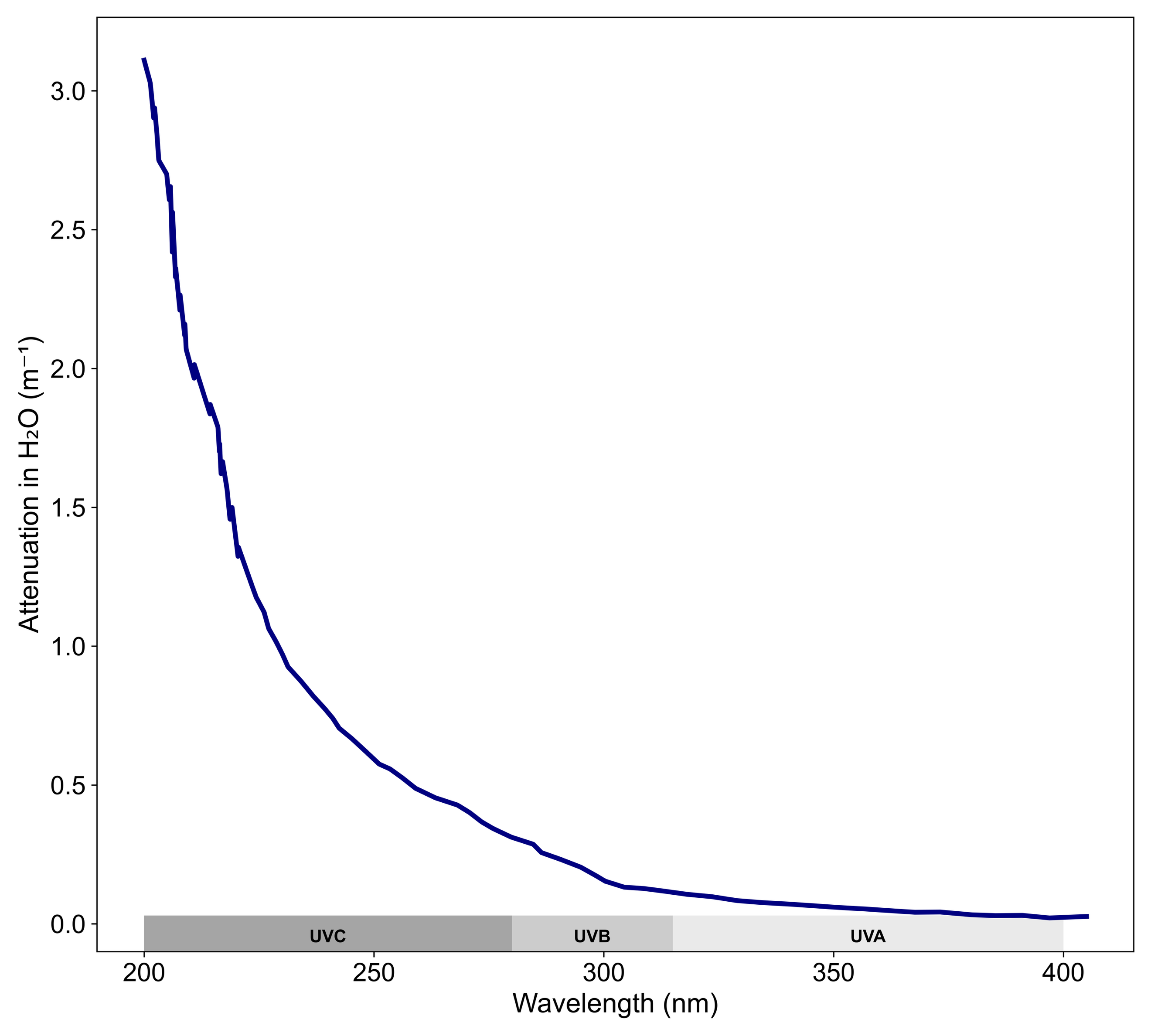

**Figure S11.** Absorption spectrum for water from Hale & Querry (1973) used to generate the irradiance spectra in the ocean’s mixed layer.

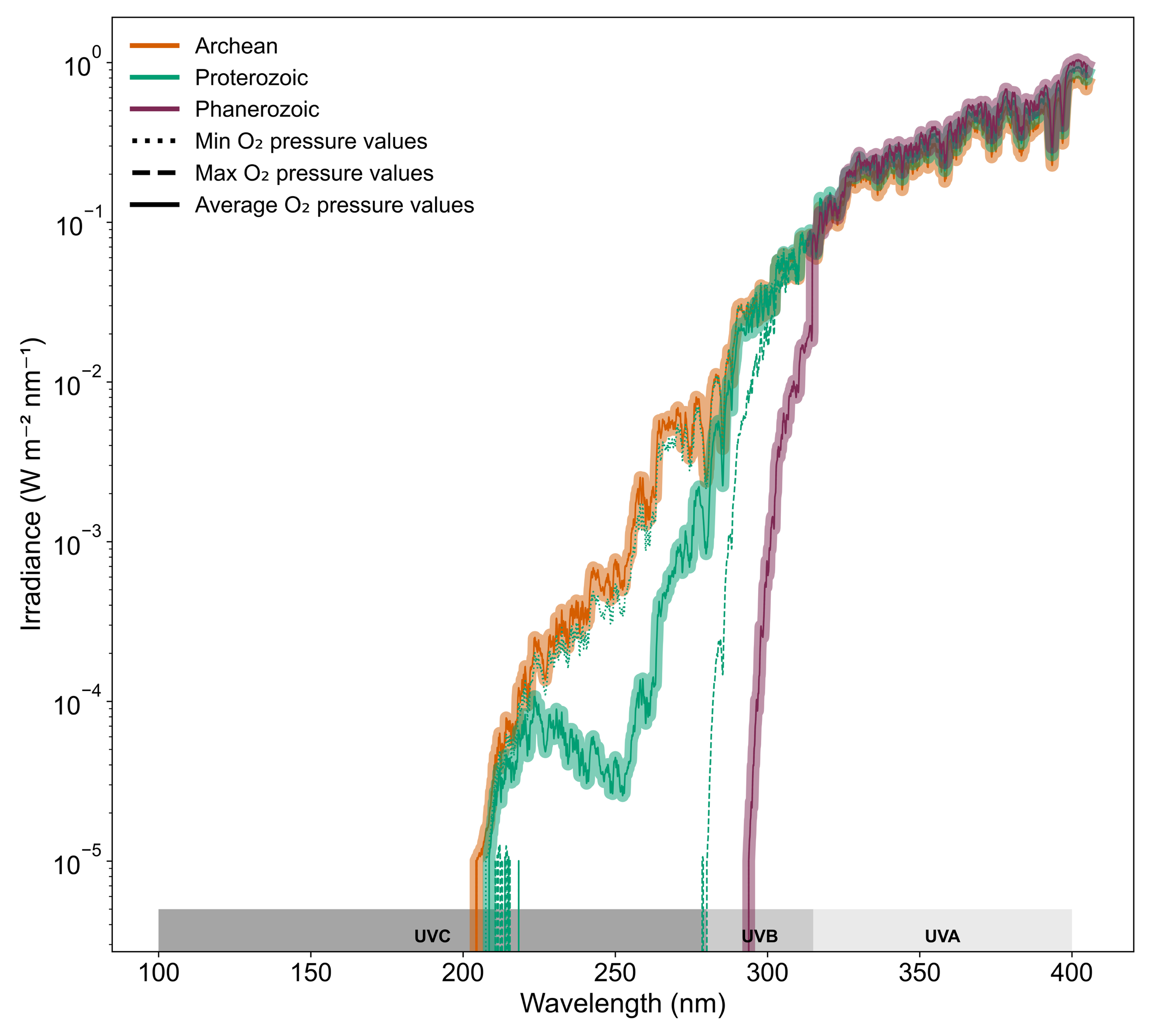

**Figure S12.** Irradiance spectra in the ocean mixed layer for the Archean, Proterozoic, and Phanerozoic. The mixed layer was set to a depth of 50 meters. The spectra were obtained by applying water’s absorption coefficients to the irradiance spectra at Earth’s surface using the Lambert-Beer law

**SUPPLEMENTAL TABLES**

**Table S1.** Summary of the 3,076 genomes compiled in this study. Genomes were acquired from the JGI genomic catalog of Earth’s microbiomes (GEMs) and the NCBI database. Metabolic guild was constrained by the presence of one or more marker genes.

| **Metabolic guild** | **Method of dataset construction** | **Number of genomes** |
| --- | --- | --- |
| Aerobic heterotrophy | Genomes from Gralka et al. (2023) and genomes containing cytochrome c oxidase (*COX2*) marker gene | 603 |
| Fermentation | Genomes from the database compiled by Zhang & Hackmann (2023) that were not from engineered environments | 198 |
| Dissimilatory ferric iron reduction | Marker genes integral outer-membrane β-barrel protein (*mtrB)*, periplasmic c-type cytochrome (*mtrA),* and tetraheme cytochrome c *(cymA)* | 378 |
| Dissimilatory nitrate reduction | Marker gene dissimilatory nitrate reductase subunit alpha (*narG)* | 483 |
| Dissimilatory sulfate reduction | Marker gene dissimilatory sulfite reductase subunit alpha (*dsrA)* | 476 |
| Methanogenesis | Marker gene methyl coenzyme M reductase (*mcrA)* | 335 |
| Anoxygenic phototrophy | Marker gene bacteriochlorophyll synthesis pathway protein (*bchC*) | 274 |
| Oxygenic phototrophy | Genomes of Cyanobacteriota | 329 |

**Table S2.** List and sources of hidden Markov models (HMMs) of marker genes used to classify genomes into distinct metabolic guilds.

| **Metabolic guild** | **Marker gene** | **HMM Accession / Source** | **Bit score cutoff** |
| --- | --- | --- | --- |
| Aerobic heterotrophy | *COX2* | Appler et al. 2025 | *e-value 1x10^-50^* |
| Dissimilatory ferric iron reduction | *mtrB* | TIGR03509.1 | 240.1 |
|  | *mtrA* | Garber et al. 2020 | 140 |
|  | *cymA* | Garber et al. 2020 | 237 |
| Dissimilatory nitrate reduction | *narG* | TIGR01580.1 | 1084.2 |
| Dissimilatory sulfate reduction | *dsrA* | TIGR02064.1 | 392.3 |
| Methanogenesis | *mcrA* | TIGR03256.1 | 767.8 |
| Anoxygenic phototrophy | *bchC* | TIGR01202.1 | 291.3 |

**Table S3.** Hydrolytic enzyme types investigated in this study, based on their classification according to the International Union of Biochemistry and Molecular Biology (IUBMB), their functions and their source. The enzyme names used to search microbial genomes are underlined.

| **Enzyme** | | | **Function** | **Substrate** | **Source** | **Refs** |
| --- | --- | --- | --- | --- | --- | --- |
| **Carbohydrate-acting** | | | |  |  |  |
|  | Glycosyl hydrolase (EC 3.2.1)  (Glycosidase, Glycoside hydrolase) | | Catalyze the hydrolysis of glycosidic bonds | Carbohydrate-containing | All domains | 1,2 |
|  |  | Alginase | Catalyze the hydrolysis of algin | Algin (*β*-1,4-D-mannuronate;  *α*-1,4-L-guluronate) | Brown algae | 2,3,4 |
|  |  | Amylase | Catalyze the hydrolysis of starch and glycogen | Starch and glycogen  (*α*-1,4- ;*α*-1,6-glucans, differ in their degree of branching) | Plants (starch)  Animals, bacteria, and yeasts (glycogen) | 2,3 |
|  |  | Arabinosidase | Catalyze the hydrolysis of arabinoside | Arabinose-containing | Plants and bacteria | 2 |
|  |  | Cellulase | Catalyze the hydrolysis of cellulose | Cellulose (*β*-1,4-glucan) | Plants and tunicates (animals) | 2 |
|  |  | Cellobiosidase | Catalyze the hydrolysis of cellulose, releasing cellobiose |  |  |  |
|  |  | Chitinase | Catalyze the hydrolysis of chitin | Chitin (*β*‐1,4‐linked *N*‐acetyl‐D‐glucosamine) | Arthropods, fungi, annelids and some plants | 2,3 |
|  |  | Fructosidase | Catalyze the hydrolysis of a fructoside | Fructose-containing | All domains | 5 |
|  |  | Furanosidase | Catalyze the hydrolysis of a furanoside | Furanose-containing | All domains | 6 |
|  |  | Glucosidase (Glucohydrolase) | Catalyze the hydrolysis of glycosidic bonds, releasing glucose | Glucose-containing | All domains | 1,2 |
|  |  | Glucosaminidase | Catalyze the hydrolysis of chitin, releasing glucosamine | Chitin (*β*‐1,4‐linked *N*‐acetyl‐D‐glucosamine) | Arthropods, fungi, annelids and some plants | 1,2,3 |
|  |  | Laminarinase | Catalyze the hydrolysis of laminarin | Laminarin (*β*-1,3-glucan) | Brown algae (*Laminaria* spp.) | 2,3 |
|  |  | Pullulanase | Catalyze the hydrolysis of pullulan | Starch-derived linear polysaccharide (*α*-1,4- ;*α*-1,6-glucan) | Fungi | 2 |
|  |  | Xylanase | Catalyze the hydrolysis of xylan | Xylan (*β*-1,4-xylose) | Plants and green algae | 2,3 |
|  |  | Xylosidase | Catalyze the hydrolysis of xylose | Xylose-containing | Plants and green algae | 2 |
|  | Carbohydrate esterase | | Cleave ‘accessory’ groups on polysaccharides | Carbohydrate-containing | All domains | 1 |
| **Protein-acting** | | |  |  |  |  |
|  | Peptidase (Protease, Proteinase) (EC 3.2.4) | | Catalyze the hydrolysis of peptide bonds | Proteins | All domains | 1 |

EC = Enzyme Commission number.

^1^McDonald et al. 2009; ^2^Oxford Dictionary of Biochemistry and Molecular Biology; ^3^Vonk & Western, 1984; ^4^Janeček & Svensson, 2022; ^5^ Kırtel et al. 2019; ^6^Naumoff, 2011

**Table S4.** Annotations of genes containing signal peptide sequences with extracellular localization that were omitted from quantification of exoenzymes. These genes corresponded to enzymes involved in housekeeping or virulence activity.

**Gene annotation**

Beta-barrel_assembly-enhancing_protease_translation

D-Ala-D-Ala_carboxypeptidase_3_(S13)_family_protein_translation

D-alanyl-D-alanine_carboxypeptidase_DacC_translation

Gamma-D-glutamyl-L-lysine_dipeptidyl-peptidase_translation

Glycyl-glycine_endopeptidase_LytM_translation

Immune_inhibitor_A_peptidase_M6_translation

Immunoglobulin_A1_protease_autotransporter_translation

Immunoglobulin_A1_protease_translation

Immunomodulating_metalloprotease_translation

L,D-transpeptidase_5_translation

L,D-transpeptidase_catalytic_domain_translation

Lysyl_endopeptidase_translation

Metalloprotease_StcE_translation

Metalloprotease_YcaL_translation

Minor_extracellular_protease_Epr_translation

Minor_extracellular_protease_vpr_translation

Peptidoglycan_DL-endopeptidase_CwlO_translation

Peptidoglycan_endopeptidase_RipA_translation

Peptidoglycan_endopeptidase_RipB_translation

Pre-pro-metalloprotease_PrtV_translation

Protease_inhibitor_translation

Serine_protease_pic_autotransporter_translation

Transglutaminase-activating_metalloprotease_translation

Tricorn_protease_translation

Virulence_metalloprotease_translation

Zinc_metalloproteinase_aureolysin_translation

**Table S5.** List and sources of Hidden Markov Models (HMMs) used to find homologs of alkaline phosphatase related genes

| **Enzyme** | **Marker gene** | **HMM accession** | **E-value threshold** |
| --- | --- | --- | --- |
| Alkaline phosphatase | *phoA* | NF007810.0 | 0.001 |
| Alkaline phosphatase | *phoD* | PF09423.14 | 0.001 |
| Alkaline phosphatase | *phoX* | PF05787.17 | 0.001 |

**Table S6.** Estimated dates (in millions of years ago) for the spread of genes encoding extracellular PhoD into new microbial lineages via speciation (spe), horizontal gene transfer (hgt), and duplication (dup). Gene events are shown under the default event cost of HGT = 3 in ecceTERA. Asterisks (*) are next to median / midpoint indicate gene events that occur at the same time when estimated with the same clock models and different event costs.

|  | CIR | | | LN | | | UGAM | | |
| --- | --- | --- | --- | --- | --- | --- | --- | --- | --- |
| Event type | Median / Midpoint | Oldest CI | Youngest CI | Median / Midpoint | Oldest CI | Youngest CI | Median / Midpoint | Oldest CI | Youngest CI |
| spe | **2,695*** | 2,864 | 2,512 | **1,958*** | 2,196 | 1,710 | **1,969** | 2,330 | 1,622 |
| spe | **2,491*** | 2,692 | 2,258 | **1,662*** | 1,920 | 1,393 | **1,598** | 1,979 | 1,216 |
| spe | **1,392*** | 1,498 | 1,259 | **1,305*** | 1,417 | 1,172 | **1,337** | 1,587 | 310 |
| hgt | **1,325*** | 2,803 | 0 | **967*** | 2,142 | 0 | **930** | 2,172 | 0 |
| hgt | **1,255*** | 2,647 | 0 | **944*** | 2,062 | 0 | **930*** | 2,237 | 0 |
| dup | **1,040*** | 2,310 | 0 | **606*** | 1,444 | 0 | **644** | 1,637 | 0 |
| hgt | **1,040*** | 2,310 | 0 | **606*** | 1,444 | 0 | **644** | 1,637 | 0 |
| dup | **1,022*** | 2,149 | 0 | **939*** | 1,990 | 0 | **868*** | 1,915 | 0 |
| hgt | **1,010** | 2,176 | 0 | **666** | 1,490 | 0 | **797*** | 1,776 | 0 |
| hgt | **987** | 2,139 | 0 | **636** | 1,421 | 0 | **724*** | 1,661 | 0 |
| spe | **941*** | 1,164 | 671 | **873*** | 1,068 | 688 | **959** | 1,459 | 154 |
| hgt | **837*** | 1,901 | 0 | **484*** | 1,155 | 0 | **594*** | 1,426 | 0 |
| hgt | **573*** | 1,396 | 0 | **440*** | 1,126 | 0 | **381** | 1,388 | 0 |

CIR = Cox-Ingersoll-Ross; LN = lognormal; UGAM = uncorrelated gamma multipliers; CI = confidence interval

**Table S7.** Estimated dates (in millions of years ago) for the spread of genes encoding extracellular PhoD into new microbial lineages via speciation (spe), horizontal gene transfer (hgt), and duplication (dup). Gene events are shown under the modified event cost of HGT = 2 in ecceTERA. Asterisks (*) are next to median / midpoint indicate gene events that occur at the same time when estimated with the same clock models and different event costs.

|  | CIR | | | LN | | | UGAM | | |
| --- | --- | --- | --- | --- | --- | --- | --- | --- | --- |
| Event type | Median / Midpoint | Oldest CI | Youngest CI | Median / Midpoint | Oldest CI | Youngest CI | Median / Midpoint | Oldest CI | Youngest CI |
| spe | **2,695*** | 2,864 | 2,512 | **1,958*** | 2,196 | 1,710 | **n/a** | n/a | n/a |
| spe | **2,491*** | 2,692 | 2,258 | **1,662*** | 1,920 | 1,393 | **n/a** | n/a | n/a |
| spe | **1,392*** | 1,498 | 1,259 | **1,305*** | 1,417 | 1,172 | **n/a** | n/a | n/a |
| hgt | **1,325*** | 2,803 | 0 | **967*** | 2,142 | 0 | **724*** | 1,661 | 0 |
| hgt | **1,255*** | 2,647 | 0 | **944*** | 2,062 | 0 | **797** | 1,776 | 0 |
| hgt | **1,245** | 2,692 | 0 | **831** | 1,920 | 0 | **930** | 2,237 | 0 |
| hgt | **1,040*** | 2,310 | 0 | **606*** | 1,444 | 0 | **797*** | 1,776 | 0 |
| hgt | **1,040*** | 2,310 | 0 | **606*** | 1,444 | 0 | **594** | 1,426 | 0 |
| dup | **1,022*** | 2,149 | 0 | **939*** | 1,990 | 0 | **868** | 1,915 | 0 |
| hgt | **1,022** | 2,149 | 0 | **939** | 1,990 | 0 | **930*** | 2,237 | 0 |
| hgt | **987** | 2,139 | 0 | **636** | 1,421 | 0 | **724** | 1,661 | 0 |
| spe | **941*** | 1,164 | 671 | **873*** | 1,068 | 688 | **n/a** | n/a | n/a |
| hgt | **837*** | 1,901 | 0 | **484*** | 1,155 | 0 | **594** | 1,426 | 0 |
| hgt | **573*** | 1,396 | 0 | **440*** | 1,126 | 0 | **930** | 2,237 | 0 |
| dup | **n/a** | n/a | n/a | **n/a** | n/a | n/a | **868*** | 1,915 | 0 |
| hgt | **n/a** | n/a | n/a | **n/a** | n/a | n/a | **797** | 1,776 | 0 |
| hgt | **n/a** | n/a | n/a | **n/a** | n/a | n/a | **594*** | 1,426 | 0 |
| hgt | **n/a** | n/a | n/a | **n/a** | n/a | n/a | **724** | 1,661 | 0 |
| dup | **n/a** | n/a | n/a | **n/a** | n/a | n/a | **868** | 1,915 | 0 |

CIR = Cox-Ingersoll-Ross; LN = lognormal; UGAM = uncorrelated gamma multipliers; CI = confidence interval

**Table S8.** Estimated dates (in millions of years ago) for the spread of genes encoding extracellular PhoD into new microbial lineages via speciation (spe), horizontal gene transfer (hgt), and duplication (dup). Gene events are shown under the modified event cost of HGT = 4 in ecceTERA. Asterisks (*) are next to median / midpoint indicate gene events that occur at the same time when estimated with the same clock models and different event costs.

|  | CIR | | | LN | | | UGAM | | |
| --- | --- | --- | --- | --- | --- | --- | --- | --- | --- |
| Event type | Median / Midpoint | Oldest CI | Youngest CI | Median / Midpoint | Oldest CI | Youngest CI | Median / Midpoint | Oldest CI | Youngest CI |
| spe | **2,695*** | 2,864 | 2,512 | **1,958*** | 2,196 | 1,710 | **n/a** | n/a | n/a |
| spe | **2,491*** | 2,692 | 2,258 | **1,662*** | 1,920 | 1,393 | **n/a** | n/a | n/a |
| spe | **1,392*** | 1,498 | 1,259 | **1,305*** | 1,417 | 1,172 | **n/a** | n/a | n/a |
| hgt | **1,325*** | 2,803 | 0 | **967*** | 2,142 | 0 | **797*** | 1,776 | 0 |
| hgt | **1,255*** | 2,647 | 0 | **944*** | 2,062 | 0 | **930*** | 2,237 | 0 |
| dup | **1,040*** | 2,310 | 0 | **606*** | 1,444 | 0 | **868*** | 1,915 | 0 |
| hgt | **1,040*** | 2,310 | 0 | **606*** | 1,444 | 0 | **930** | 2,237 | 0 |
| dup | **1,022*** | 2,149 | 0 | **939*** | 1,990 | 0 | **868** | 1,915 | 0 |
| hgt | **1,010** | 2,176 | 0 | **666** | 1,490 | 0 | **797** | 1,776 | 0 |
| hgt | **987** | 2,139 | 0 | **636** | 1,421 | 0 | **724** | 1,661 | 0 |
| spe | **941*** | 1,164 | 671 | **873*** | 1,068 | 688 | **n/a** | n/a | n/a |
| hgt | **837*** | 1,901 | 0 | **484*** | 1,155 | 0 | **594** | 1,426 | 0 |
| hgt | **573*** | 1,396 | 0 | **440*** | 1,126 | 0 | **930** | 2,237 | 0 |
| hgt | **n/a** | n/a | n/a | **n/a** | n/a | n/a | **594** | 1,426 | 0 |
| hgt | **n/a** | n/a | n/a | **n/a** | n/a | n/a | **724*** | 1,661 | 0 |
| hgt | **n/a** | n/a | n/a | **n/a** | n/a | n/a | **797** | 1,776 | 0 |
| hgt | **n/a** | n/a | n/a | **n/a** | n/a | n/a | **594*** | 1,426 | 0 |
| hgt | **n/a** | n/a | n/a | **n/a** | n/a | n/a | **724** | 1,661 | 0 |
| dup | **n/a** | n/a | n/a | **n/a** | n/a | n/a | **868** | 1,915 | 0 |

CIR = Cox-Ingersoll-Ross; LN = lognormal; UGAM = uncorrelated gamma multipliers; CI = confidence interval

**Table S9.** Estimated dates (in millions of years ago) for the spread of genes encoding extracellular PhoD into new microbial lineages via speciation (spe), horizontal gene transfer (hgt), and duplication (dup). Gene events are shown under the modified event cost of HGT = 6 in ecceTERA. Asterisks (*) are next to median / midpoint indicate gene events that occur at the same time when estimated with the same clock models and different event costs.

|  | CIR | | | LN | | | UGAM | | |
| --- | --- | --- | --- | --- | --- | --- | --- | --- | --- |
| Event type | Median / Midpoint | Oldest CI | Youngest CI | Median / Midpoint | Oldest CI | Youngest CI | Median / Midpoint | Oldest CI | Youngest CI |
| spe | **2,695*** | 2,864 | 2,512 | **1,958*** | 2,196 | 1,710 | **n/a** | n/a | n/a |
| spe | **2,491*** | 2,692 | 2,258 | **1,662*** | 1,920 | 1,393 | **n/a** | n/a | n/a |
| spe | **2,287** | 2,418 | 2,141 | **1,643** | 1,782 | 1,503 | **n/a** | n/a | n/a |
| spe | **2,227** | 2,362 | 2,074 | **1,566** | 1,704 | 1,426 | **n/a** | n/a | n/a |
| spe | **2,185** | 2,327 | 2,027 | **1,516** | 1,663 | 1,364 | **n/a** | n/a | n/a |
| spe | **2,136** | 2,279 | 1,976 | **1,454** | 1,595 | 1,308 | **n/a** | n/a | n/a |
| spe | **2,020** | 2,176 | 1,841 | **1,332** | 1,490 | 1,168 | **n/a** | n/a | n/a |
| spe | **1,974** | 2,139 | 1,788 | **1,273** | 1,421 | 1,120 | **n/a** | n/a | n/a |
| spe | **1,392*** | 1,498 | 1,259 | **1,305*** | 1,417 | 1,172 | **n/a** | n/a | n/a |
| hgt | **1,325*** | 2,803 | 0 | **967*** | 2,142 | 0 | **797*** | 1,776 | 0 |
| hgt | **1,255*** | 2,647 | 0 | **944*** | 2,062 | 0 | **797** | 1,776 | 0 |
| dup | **1,040*** | 2,310 | 0 | **606*** | 1,444 | 0 | **868*** | 1,915 | 0 |
| hgt | **1,040*** | 2,310 | 0 | **606*** | 1,444 | 0 | **930** | 2,237 | 0 |
| dup | **1,022*** | 2,149 | 0 | **939*** | 1,990 | 0 | **868** | 1,915 | 0 |
| hgt | **1,022** | 2,149 | 0 | **939** | 1,990 | 0 | **930*** | 2,237 | 0 |
| spe | **941*** | 1,164 | 671 | **873*** | 1,068 | 688 | **n/a** | n/a | n/a |
| hgt | **837*** | 1,901 | 0 | **484*** | 1,155 | 0 | **594** | 1,426 | 0 |
| hgt | **573*** | 1,396 | 0 | **440*** | 1,126 | 0 | **930** | 2,237 | 0 |
| hgt | **n/a** | n/a | n/a | **n/a** | n/a | n/a | **724** | 1,661 | 0 |
| hgt | **n/a** | n/a | n/a | **n/a** | n/a | n/a | **594** | 1,426 | 0 |
| hgt | **n/a** | n/a | n/a | **n/a** | n/a | n/a | **724*** | 1,661 | 0 |
| hgt | **n/a** | n/a | n/a | **n/a** | n/a | n/a | **797** | 1,776 | 0 |
| hgt | **n/a** | n/a | n/a | **n/a** | n/a | n/a | **594*** | 1,426 | 0 |
| hgt | **n/a** | n/a | n/a | **n/a** | n/a | n/a | **724** | 1,661 | 0 |
| dup | **n/a** | n/a | n/a | **n/a** | n/a | n/a | **868** | 1,915 | 0 |

CIR = Cox-Ingersoll-Ross; LN = lognormal; UGAM = uncorrelated gamma multipliers; CI = confidence interval

**References**

Appler, K. E. et al. Oxygen metabolism in descendants of the archaeal-eukaryotic ancestor. Preprint at bioRxiv https://doi.org/10.1101/2024.07.04.601786v1 (2025)

Altschul, S. Gapped BLAST and PSI-BLAST: a new generation of protein database search programs. *Nucleic Acids Research* **25**, 3389–3402 (1997).

Bianchini, G. & Sánchez‐Baracaldo, P. TreeViewer : Flexible, modular software to visualise and manipulate phylogenetic trees. *Ecology and Evolution* **14**, e10873 (2024).

Boden, J. S., Zhong, J., Anderson, R. E. & Stüeken, E. E. Timing the evolution of phosphorus-cycling enzymes through geological time using phylogenomics. *Nat Commun* **15**, 3703 (2024).

Bowers, R. *et al.* Minimum information about a single amplified genome (MISAG) and a metagenome-assembled genome (MIMAG) of bacteria and archaea. *Nat Biotechnol* **35**, 725–731 (2017).

Capella-Gutiérrez, S., Silla-Martínez, J. M. & Gabaldón, T. trimAl: a tool for automated alignment trimming in large-scale phylogenetic analyses. *Bioinformatics* **25**, 1972–1973 (2009).

Cavalazzi, B. *et al.* Cellular remains in a ~3.42-billion-year-old subseafloor hydrothermal environment. *Sci. Adv.* **7**, eabf3963 (2021).

Claire, M., Sheets, J., Cohen, M., Ribas, I., Meadows, V., & Catling, D. (2012). The Evolution of Solar Flux from 0.1nm to 160 μm: Quantitative Estimates for Planetary Studies. *The Astrophysical Journal*, *757*(1), 95–106.

Cnossen, I. *et al.* Habitat of early life: Solar X‐ray and UV radiation at Earth’s surface 4–3.5 billion years ago. *J. Geophys. Res.* **112**, 2006JE002784 (2007).

. Cooke, G. J., Marsh, D. R., Walsh, C., Black, B. & Lamarque, J.-F. A revised lower estimate of ozone columns during Earth’s oxygenated history. *R. Soc. open sci.* **9**, 211165 (2022).

David, L. A. & Alm, E. J. Rapid evolutionary innovation during an Archaean genetic expansion. *Nature* **469**, 93–96 (2011).

Eddy, S. R. Accelerated Profile HMM Searches. *PLoS Comput Biol* **7**, e1002195 (2011).

Fakhraee, M., Tarhan, L. G., Planavsky, N. J. & Reinhard, C. T. A largely invariant marine dissolved organic carbon reservoir across Earth’s history. *Proc Natl Acad Sci U S A* **118**, e2103511118 (2021).

Garber, A. I. *et al.* FeGenie: A Comprehensive Tool for the Identification of Iron Genes and Iron Gene Neighborhoods in Genome and Metagenome Assemblies. *Front. Microbiol.* **11**, 37 (2020).

Gralka, M., Pollak, S. & Cordero, O. X. Genome content predicts the carbon catabolic preferences of heterotrophic bacteria. *Nat Microbiol* **8**, 1799–1808 (2023).

Green, E. R. & Mecsas, J. Bacterial Secretion Systems: An Overview. *Microbiol Spectr* **4**, 4.1.13 (2016).

Hackmann, T. J. & Zhang, B. The phenotype and genotype of fermentative prokaryotes. *Sci. Adv.* **9**, eadg8687 (2023).

Hale, G. M. & Querry, M. R. Optical Constants of Water in the 200-nm to 200-μm Wavelength Region. *Appl. Opt.* **12**, 555 (1973).

Hoang, D. T., Chernomor, O., Von Haeseler, A., Minh, B. Q. & Vinh, L. S. UFBoot2: Improving the Ultrafast Bootstrap Approximation. *Molecular Biology and Evolution* **35**, 518–522 (2018).Jacox, E., Chauve, C., Szöllősi, G. J., Ponty, Y. & Scornavacca, C. ecceTERA: comprehensive gene tree-species tree reconciliation using parsimony. *Bioinformatics* **32**, 2056–2058 (2016).

Janeček, Š. & Svensson, B. How many α-amylase GH families are there in the CAZy database? *Amylase* **6**, 1–10 (2022).

Kalyaanamoorthy, S., Minh, B. Q., Wong, T. K. F., Von Haeseler, A. & Jermiin, L. S. ModelFinder: fast model selection for accurate phylogenetic estimates. *Nat Methods* **14**, 587–589 (2017).

Katoh, K. & Standley, D. M. MAFFT Multiple Sequence Alignment Software Version 7: Improvements in Performance and Usability. *Molecular Biology and Evolution* **30**, 772–780 (2013).

Kaushik, S., He, H. & Dalbey, R. E. Bacterial Signal Peptides- Navigating the Journey of Proteins. *Front. Physiol.* **13**, (2022).

Kırtel, O., Lescrinier, E., Van Den Ende, W. & Toksoy Öner, E. Discovery of fructans in Archaea. *Carbohydrate Polymers* **220**, 149–156 (2019).

Lartillot, N., Lepage, T. & Blanquart, S. PhyloBayes 3: a Bayesian software package for phylogenetic reconstruction and molecular dating. *Bioinformatics* **25**, 2286–2288 (2009). 1.

Lehmer, O. R., Catling, D. C., Buick, R., Brownlee, D. E. & Newport, S. Atmospheric CO_2_ levels from 2.7 billion years ago inferred from micrometeorite oxidation. *Sci. Adv.* **6**, eaay4644 (2020).

Lyons, T., Reinhard, C., & Planavsky, N. (2014). The rise of oxygen in Earth’s early ocean and atmosphere. *Nature*, *506*, 307–315.

Mateos, K. *et al.* The evolution and spread of sulfur cycling enzymes reflect the redox state of the early Earth. *Science Advances* **9**, eade4847 (2023).

McDonald, A. G., Boyce, S. & Tipton, K. F. ExplorEnz: the primary source of the IUBMB enzyme list. *Nucleic Acids Research* **37**, D593–D597 (2009).

Minh, B. Q. *et al.* IQ-TREE 2: New Models and Efficient Methods for Phylogenetic Inference in the Genomic Era. *Molecular Biology and Evolution* **37**, 1530–1534 (2020).Moody, E. R. R. *et al.* The nature of the last universal common ancestor and its impact on the early Earth system. *Nat Ecol Evol* **8**, 1654–1666 (2024).

Nakamura, T., Yamada, K. D., Tomii, K. & Katoh, K. Parallelization of MAFFT for large-scale multiple sequence alignments. *Bioinformatics* **34**, 2490–2492 (2018).

Naumoff, D. G. Hierarchical classification of glycoside hydrolases. *Biochemistry Moscow* **76**, 622–635 (2011).

Owji, H., Nezafat, N., Negahdaripour, M., Hajiebrahimi, A. & Ghasemi, Y. A comprehensive review of signal peptides: Structure, roles, and applications. *European Journal of Cell Biology* **97**, 422–441 (2018).

*Oxford Dictionary of Biochemistry and Molecular Biology*. (Oxford University Press, Oxford ; New York, 2006).

Papanikou, E., Karamanou, S. & Economou, A. Bacterial protein secretion through the translocase nanomachine. *Nat Rev Microbiol* **5**, 839–851 (2007).

Parks, D. H., Imelfort, M., Skennerton, C. T., Hugenholtz, P. & Tyson, G. W. CheckM: assessing the quality of microbial genomes recovered from isolates, single cells, and metagenomes. *Genome Res.* **25**, 1043–1055 (2015).

Pugsley, A. P., Francetic, O., Driessen, A. J. & Lorenzo, V. D. Getting out: protein traffic in prokaryotes. *Molecular Microbiology* **52**, 3–11 (2004).

Robinson, C. & Bolhuis, A. Tat-dependent protein targeting in prokaryotes and chloroplasts. *Biochimica et Biophysica Acta (BBA) - Molecular Cell Research* **1694**, 135–147 (2004).

Ronquist, F., Teslenko, M., Van Der Mark, P., Ayres, D. L., Darling, A., Hohna, S., Larget, B., Liu, L., Suchard, M. A. & Huelsenbeck, J. P. MrBayes 3.2: Efficient Bayesian phylogenetic inference and model choice across a large model space. *Systematic Biology,* **61,** 539-542 (2012).

Seemann, T. Prokka: rapid prokaryotic genome annotation. *Bioinformatics* **30**, 2068–2069 (2014).

Shannon, P. *et al.* Cytoscape: A Software Environment for Integrated Models of Biomolecular Interaction Networks. *Genome Res.* **13**, 2498–2504 (2003).

Shimodaira, H. An Approximately Unbiased Test of Phylogenetic Tree Selection. *Systematic Biology* **51**, 492–508 (2002).

Som, S. M. *et al.* Earth’s air pressure 2.7 billion years ago constrained to less than half of modern levels. *Nature Geosci* **9**, 448–451 (2016).Teufel, F. *et al.* SignalP 6.0 predicts all five types of signal peptides using protein language models. *Nat Biotechnol* **40**, 1023–1025 (2022).

Tully, B. J., Graham, E. D. & Heidelberg, J. F. The reconstruction of 2,631 draft metagenome-assembled genomes from the global oceans. *Sci Data* **5**, 170203 (2018).

Ueno, Y., Yamada, K., Yoshida, N., Maruyama, S. & Isozaki, Y. Evidence from fluid inclusions for microbial methanogenesis in the early Archaean era. *Nature* **440**, 516–519 (2006).

Vonk, H. J. & Western, J. R. H. *Comparative Biochemistry and Physiology of Enzymatic Digestion*. (Acad. Press, London, 1984).

Witwinowski, J. *et al.* An ancient divide in outer membrane tethering systems in bacteria suggests a mechanism for the diderm-to-monoderm transition. *Nat Microbiol* **7**, 411–422 (2022).

Wolfe, J. M. & Fournier, G. P. Horizontal gene transfer constrains the timing of methanogen evolution. *Nat Ecol Evol* **2**, 897–903 (2018).

Wu, L. F., Ize, B., Chanal, A., Quentin, Y. & Fichant, G. Bacterial twin-arginine signal peptide-dependent protein translocation pathway: evolution and mechanism. *J Mol Microbiol Biotechnol* **2**, 179–189 (2000).

Yu, N. Y. *et al.* PSORTb 3.0: improved protein subcellular localization prediction with refined localization subcategories and predictive capabilities for all prokaryotes. *Bioinformatics* **26**, 1608–1615 (2010).

Zhang, C., Zhao, Y., Braun, E. L. & Mirarab, S. TAPER: Pinpointing errors in multiple sequence alignments despite varying rates of evolution. *Methods in Ecology and Evolution,* **12,** 2145-2158 (2021).
